# Capabilities, specificity gaps and training-data dependence of AlphaFold3 across diverse application areas

**DOI:** 10.64898/2026.07.13.738147

**Authors:** Océane Follonier, Yan Liu, Pablo Campomanes, Luc Lafrenaye, Julien Racle, Daniel Alvarez, Julian van Gerwen, Ricardo Heinzmann, Jürgen Jänes, Eric Kummelstedt, Janani Durairaj, David Gfeller, Stefano Vanni, Pedro Beltrao

## Abstract

Structure prediction models have moved from single proteins to assemblies that include diverse biomolecules and their modifications. AlphaFold3 (AF3) and related models extended structural modelling via an all-atom framework, opening many new potential applications in structural biology. We evaluate how well the new capabilities of AF3 translate into application tasks in diverse areas: prediction of ubiquitinated protein structures, T-cell receptor (TCR)-epitope recognition, antibody-antigen complexes, protein-RNA and protein-lipid interactions. We find that, while AF3 can perform well in favourable settings, this performance is uneven across applications. In RNA-target predictions, the model confidence fails to separate genuine from decoy interaction partners and in several tasks accuracy depends on the presence of related complexes in the training set. Taken together, our assessment is more cautious than for AF2, whose gains in modelling monomers and complexes were clear and broadly generalisable. AF3’s extension to new biomolecule types shows less consistent performance and generalisation. AF3 can be a powerful tool for hypothesis generation and prioritisation, but its predictions and use of confidence metrics will depend strongly on the specific application area and must be interpreted with respect to training-set overlap. We expect that the benchmarks provided here will serve for testing of future developments in the structure prediction field.

## Introduction

Protein structure prediction, as a field, has been a central challenge in computational biology for half a century that has seen impressive progress over the course of the last 5 years. Decades of incremental progress, through homology modelling, fragment assembly, and physics-based refinement, were transformed by deep learning models. AlphaFold2 (AF2), released around 5 years ago, reached experimental-level accuracy for well folded domains (Jumper et al. 2021), a result later reproduced by related approaches such as RoseTTAFold (Baek et al. 2021) and ESMFold (Lin et al. 2023) and soon-after extended to the modelling of protein complexes (Evans et al. 2021). The release of predicted structures for over 200 million sequences in the AlphaFold Database (Varadi et al. 2022), has led to broad availability of high confidence prediction models at an unprecedented scale.

AF2 and related methods represented a broadly useful new capability for life science research which raised the question of the extent by which such predicted models can be used in downstream applications of different kinds. A previous study detailed a community assessment that tested the limits of application of AF2 predicted structures in common downstream application in structural biology and structural bioinformatics (Akdel et al. 2022). The results of this assessment were highly encouraging showing that the predicted models, when their confidence metrics were carefully considered, could be used in diverse applications in replacement of experimental structures.

The release of AlphaFold3 (AF3) (Abramson et al. 2024) further advanced this field, by the development of a diffusion-based generative model with an all-atom representation that can be used to predict structures of proteins together with other biomolecules (e.g. RNA, DNA, lipids, etc). Soon after, several related models and open implementations of AF3 were reported including Chai-1 (Chai Discovery et al. 2024), Boltz (Wohlwend et al. 2025; Passaro et al. 2025), Protenix (Zhang et al. 2026) and OpenFold3 (<u>openfold-3.readthedocs.io</u>). These newer methods have been primarily accessed and compared in their primary task of structure prediction. Overall, there has been modest improvement in the prediction of structures of monomers and protein-protein complexes compared to AF2 with the prediction for other biomolecules having lower accuracy (Abramson et al. 2024). For antibody model predictions AF3 outperforms previous methods but it still fails on many targets, with multi-seed sampling only partially helping in identification of a correct model (Fromm et al. 2026). For protein-ligand structure predictions current co-folding methods, including AF3, seem to recover many bound poses by reproducing examples seen during training. Careful evaluation on complexes released after the training cut-off suggest a loss in accuracy for out-of-distribution ligands (Škrinjar et al. 2025).

Here, we perform a community-assessment of AF3 focusing on diverse application areas of AF3 specific capabilities, testing for practical biological applications. These areas include the prediction of ubiquitinated protein structures, TCR-epitope recognition, antibody-antigen complex prediction, protein-RNA interaction modelling, and protein-lipid co-folding. For these applications we test if AF3 can produce accurate models and/or if its confidence estimates are well calibrated, paying particular attention to two recurring questions: whether confidence discriminates genuine interactions from decoys (specificity), and whether accuracy depends on overlap with the training set. Across the application areas and evaluation approaches, the advances made by AF3 appear more nuanced than what was at the time the advances made by AF2. It is clear that AF3 expands what can be modelled but in several areas the performance seems limited in specificity and shows dependence on training-set overlap. We discuss what these results imply for the responsible use of AF3 in downstream applications.

## Results

### Prediction of ubiquitinated protein structures

Ubiquitination is a post-translational modification (PTM) in which ubiquitin is attached to lysine residues as a monomer or a polymeric chain, which can modulate the structure, interactions, and cellular fate of a target protein (Komander and Rape 2012; Yau and Rape 2016). AF3 is natively incapable of modelling protein-protein covalent bonds such as those underpinning ubiquitination (Abramson et al. 2024). However, we and others (Fábián et al. 2025) have realised that this can be circumvented by introducing a bridging small-molecule ligand, which can be covalently bound to proteins. In particular, ubiquitination can be reconstituted by converting the C-terminal glycine of ubiquitin (Gly76) into a small-molecule ligand, and specifying a peptide bond to the remainder of ubiquitin and an isopeptide bond to the ε-amino group of a target lysine residue (Fig. 1A). This opens the door for structural prediction of ubiquitination and all other forms of protein-protein covalent bonds.

**Figure 1.**
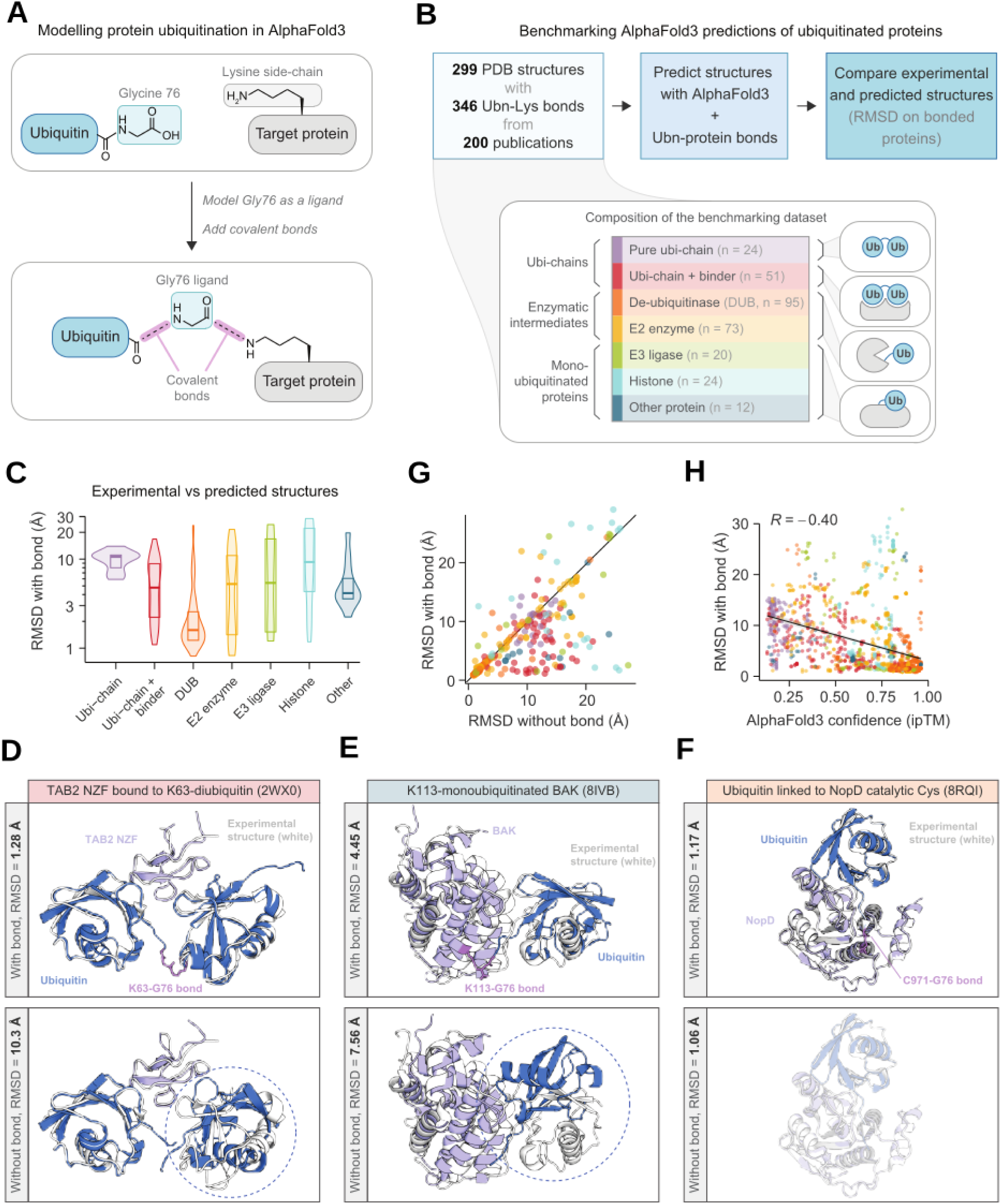
Prediction of ubiquitinated protein structures with AF3. **A)** Workflow for modelling protein ubiquitination in AlphaFold3 by converting the C-terminal glycine of ubiquitin (Gly76) into a small-molecule ligand and introducing covalent bonds. **B)** Upper: Workflow for benchmarking prediction of ubiquitinated protein structures on a dataset of experimental structures from the Protein Data Bank (PDB). Lower: Description and illustration of the types of protein structures contained in the benchmarking dataset. The number of structures in each group is shown in brackets. **C)** Root-mean-square deviation (RMSD) comparing predicted and experimental structures. RMSD was calculated on pairs of covalently bonded proteins, using the average when more than two proteins were bonded. RMSD was averaged across five AlphaFold3 models (see Methods). **D-E)** Example predicted structures of D) a ubi-chain with a binder, **E)** a mono-ubiquitinated protein, and **F)** a catalytic intermediate of a de-ubiquitinase (DUB). Predictions with and without protein-ubiquitin bonds are shown. Average RMSD values across five AlphaFold3 models are shown. Structures show a global alignment between the experimental structure (white) and a representative AlphaFold3 model (blue: ubiquitin, light blue: ubiquitinated proteins, purple: ubiquitin-protein bonds). **G)** Average RMSD across five AlphaFold3 models with protein-ubiquitin bonds included (same as C)) or excluded. H) Relationship between RMSD and AlphaFold3 model confidence (ipTM score). Protein-ubiquitin bonds were included. For each structure, the five AlphaFold3 models are shown separately. Linear regression line is shown with 95% confidence interval shaded, and Pearson’s correlation coefficient is shown.

To benchmark this approach, we constructed a database of ubiquitinated protein structures from the PDB (see Methods, Fig. 1B), containing 299 structures with 346 ubiquitin-protein bonds, including polyubiquitin chains, enzymatic intermediates of the ubiquitin-handling machinery, and mono-ubiquitinated proteins (Fig. 1B, Table S1-S2). We also included 32 structures of NEDD8 and ISG15, which structurally resemble ubiquitin but display distinct regulatory mechanisms (Table S1). Finally, 170 structures featured bonds other than Gly-Lys, including bonds to catalytic cysteine residues or non-native mimetics of ubiquitin-protein bonds (Table S2).

We used AF3 with our ligand-based approach to re-predict all 299 experimental scoring accuracy with root-mean-square deviation (RMSD) from experimental structures using only covalently bonded proteins (Table S3). Other metrics - including RMSD on the entire structure or DockQ - produced similar results (see Methods, Fig. S1). Many structures were predicted medium to high accuracy (169/299 with RMSD < 5 Å), with the greatest accuracy displayed by polyubiquitin chains interacting with non-covalent binders (median RMSD = 4.87 Å), mono-ubiquitinated non-histone proteins (median RMSD = 4.23 Å), and enzymatic intermediates of de-ubiquitinases (DUBS, median RMSD = 1.62 Å, Fig. 1c, examples in Fig. 1D-F). The lowest accuracy was seen for cryo-EM structures of mono-ubiquitinated histones - likely reflecting the size and complexity of these structures - and poly-ubiquitinated chains without binders (Fig. 1C). Because polyubiquitin chains present nearly identical inputs yet adopt diverse conformations across and within linkage types (Agrata and Komander 2025), the model likely cannot resolve the correct conformation without additional context. Similarly, AF3 failed to predict the well-studied conformational shift incurred by conjugation of NEDD8 to Cullin-5 (Fig. S2 (Duda et al. 2008)). Overall, despite variability in performance, AF3 demonstrates potential for modeling a broad range of ubiquitinated protein structures.

We next asked whether including the covalent bonds is required for accurate predictions. Running AF3 without the covalent bonds led to a widespread deterioration in accuracy (Fig. 1G), with ubiquitin placed either in the incorrect orientation or the incorrect location (examples in Fig. 1E and 1F). This leads to two conclusions: first, AF3 often cannot predict the most likely position and orientation of a ubiquitin modification from structural context alone, supporting the necessity of our workflow. Second, although most structures predate AF3 and were likely included in its training, the decline in performance observed without explicitly defining covalent bonds indicates that many high-accuracy predictions are not simply due to training data memorisation. Notably, the subset of structures that displayed high accuracy both with and without a covalent bond were dominated by enzymatic intermediates of DUBs (Fig. 1G, example in Fig. 1F), which are structurally highly similar and thus straightforward to predict.

As an additional test for generalisability, we filtered for structures where the bonded protein-ubiquitin pair was not homologous to others found in the PDB before the cutoff date for AF3 training, using a sequence identity cutoff of 30% (see Methods). While the resulting 11 structures were predicted with similar or worse accuracy compared to remaining structures of the same type (e.g. “E3 ligases”), we consider this number too small for a reliable estimate of model generalisability (Fig. S3). Finally, we examined the internal confidence metrics of AF3. Encouragingly, model confidence metrics were negatively correlated with prediction error (Fig. 1H, Fig. S4). For instance, high-confidence predictions based on the interface predicted Template Modeling (ipTM) score (ipTM > 0.8, 44% of predictions) had a median RMSD of 1.55 Å compared to 7.27 Å for remaining structures.

Here, we have demonstrated a straightforward workflow that unlocks structural prediction of ubiquitinated proteins with AF3. While we observed variable accuracy, model confidence could enrich for accurate predictions. Hence, we anticipate that this will be a useful approach for biological discovery, such as predicting binders of polyubiquitin chains, decoding structural consequences of site-specific ubiquitination, and modelling other protein-based PTMs.

### TCR-epitope interaction predictions

TCRs bind to epitopes, which consist of peptides presented on major histocompatibility complex (MHC) molecules. TCR-epitope interactions are characterized by low affinity and extensive flexibility in the TCR loops that recognize the epitopes. For this reason, standard approaches, such as homology modelling, have most often failed to predict the correct structure of TCR-epitope complexes beyond cases with extensive homology with experimental X-ray structures. A second challenge for understanding and modelling TCR-epitope interactions comes from the high diversity of MHCs (∼10^4^ alleles (Robinson et al. 2015)) and their ligands (10^10^ – 10^14^ for a given MHC (Tadros et al. 2023)) and of TCR sequences (estimates ranging from 10^16^ to 10^60^ (Murugan et al. 2012; Arstila et al. 1999)).

Prior prediction methods for TCR-epitope interactions have been mainly based on machine learning algorithms trained on large datasets of epitope-specific TCRs. These tools can reach decent accuracy for a few epitopes with abundant and reliable training data, but all of them have failed for epitopes with few or no documented TCRs (Drost et al. 2025; Nielsen et al. 2024; Richardson et al. 2026). Recent reports suggest that AlphaFold-like models can, at least in some cases, predict TCR-epitope interactions including for epitopes without known TCRs. However, prediction accuracy varies substantially across epitopes (Messemaker et al. 2025; Ascunce-París et al. 2025; Bradley 2023; Karnaukhov et al. 2024; Richardson et al. 2026; Wu et al. 2025; Woods et al. 2026; Shah et al. 2026; McMaster et al. 2026).

Here we explore different scenarios, including predictions of TCRs recognizing a given epitope out of a repertoire of TCRs, predictions of epitopes recognized by a given TCR out of a set of putative epitopes, predictions of epitope recognition by distinct TCR variants and predictions of TCR recognition for distinct epitope variants (Fig. 2A)

**Figure 2.**
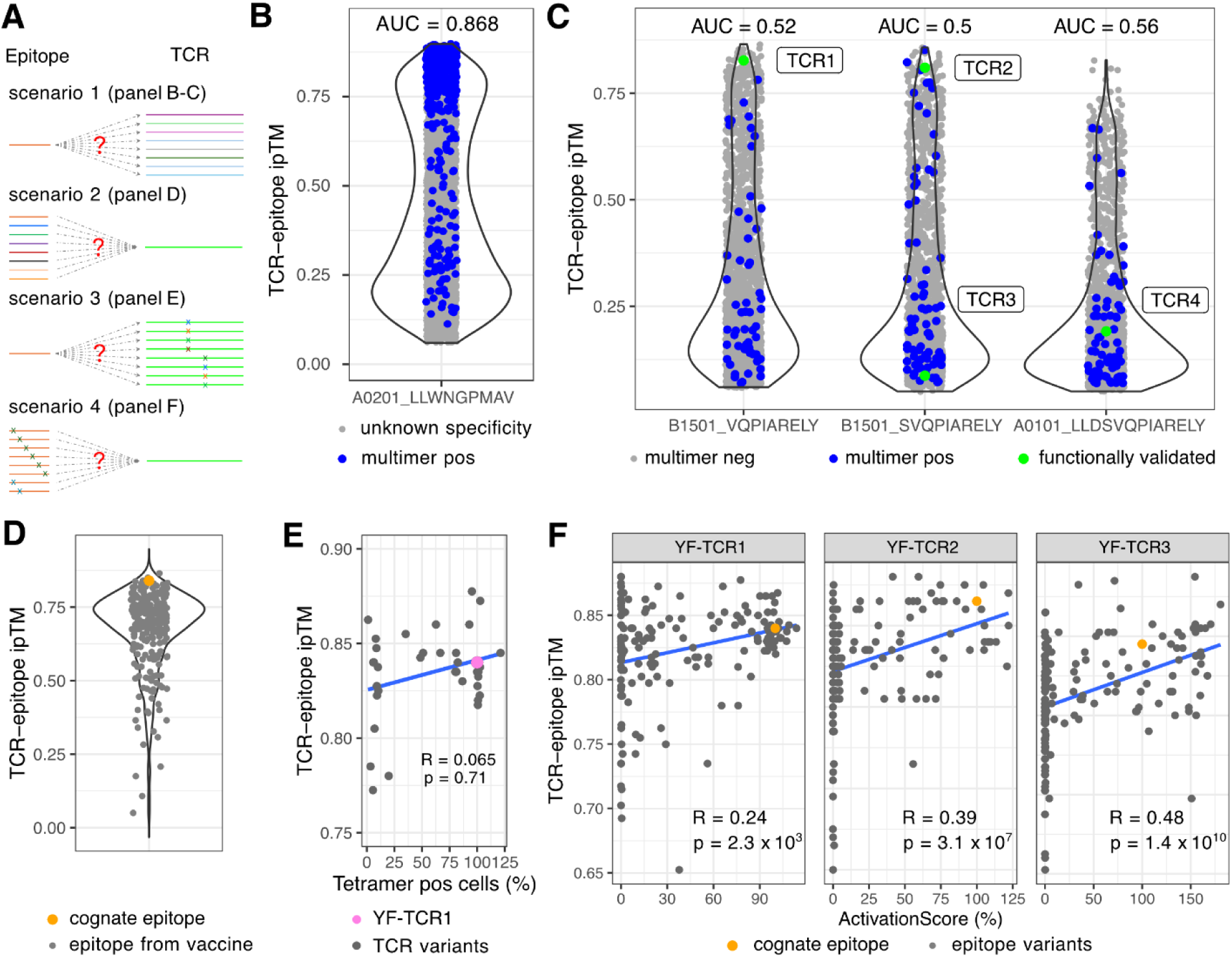
TCR-epitope interaction predictions with AF3. **A)** Schematic plots for different scenarios for benchmarking AF3 TCR-epitope interaction predictions. **B)** AF3 predictions of TCRs recognizing a viral epitope from Yellow Fever (LLWNGPMAV, restricted to HLA-A*02:01). Positives (blue) correspond to TCRs sequenced from a donor after sorting CD8 T-cells with a peptide-MHC multimer (Liu et al. 2025). Grey dots represent TCRs with unknown specificities sequenced from CD8 T cells from the same donor prior to any stimulation. **C)** Predictions of neo-antigen-specific TCRs analysed in Gumpert et al. (Gumpert et al. 2026). Positives correspond to TCRs sorted with the corresponding peptide-MHC multimers, negatives correspond to TCRs from T cells that did not bind the peptide-MHC multimers and green dots represent TCRs that were functionally validated. **D)** AF3 predictions of the recognition of HLA-A*02:01 ligands derived from YF proteome by a TCR known to interact with the LLWNGPMAV epitope (orange dot). Grey dots correspond to all other 9-mer peptides from the Yellow Fever proteome predicted to bind to HLA-A*02:01 (%rank of MixMHCpred < 2). **E)** AF3 predictions of the impact on epitope recognition of single amino acid variants in TCR sequences studied in (Liu et al. 2025). **F)** AF3 predictions of the impact on TCR recognition of all epitope single amino acid variants, based on X-scan data (Liu et al. 2025) for three TCRs (YF-TCR1, YF-TCR2 and YF-TCR3). The cognate epitope used as template in the X-scan corresponds to the YF epitope LLWNGPMAV restricted to HLA-A*02:01. Spearman correlation was used in panels E and F.

To explore whether AF3 can find TCRs recognizing a given epitope out of a repertoire of TCRs, we first leveraged data obtained from a Yellow Fever (YF) vaccinated donor for which both the baseline TCR repertoire and TCRs recognizing an immunodominant YF epitope (LLWNGPMAV, restricted to HLA-A*02:01) have been sequenced (Liu et al. 2025) (see Methods). All TCRs were modelled in complex with the epitope, and the mean AF3 ipTM between TCR chains (TCRα and TCRβ) and the epitope (peptide + MHC) was computed (Fig. 2B, Table S4A). A very clear enrichment of epitope-specific TCRs was observed among the high AF3-scoring TCRs (AUC=0.88). This demonstrates that, for this epitope for which no structural template exists in complex with a TCR, AF3 predictions are quite reliable. To expand these results in the context of cancer neo-epitope TCR recognition, we used a study where T cells recognizing three different cancer-specific neo-epitopes presented on two different MHCs had been isolated and sequenced (Gumpert et al. 2026). Four of these TCRs had been further functionally characterized. As these epitopes come from newly characterized cancer-specific mutations, no structural template exists for them in the training data of AF3. Scoring the epitope-specific TCRs with AF3 resulted in limited enrichment with respect to non-epitope-specific TCRs from the same samples (AUC < 0.6, Fig. 2C, Table S4B). However, the scores were remarkably high for two of the four functionally validated TCRs (rank 19 out of 1,946 tested TCRs for TCR1 and rank 22 out of 1,673 tested TCRs for TCR2). These two top-ranking TCRs also exhibited the strongest responses in the functional validations that were performed in the original study, while TCR3 and TCR4 only showed weak activation signals (Gumpert et al. 2026). These results demonstrate how AF3 could have been used to identify some of the TCRs recognizing both viral epitopes and cancer neo-antigens directly from the TCR repertoire of a patient.

We then investigate how AF3 could predict the epitope targeted by a TCR out of a set of peptides binding to a given MHC. Using a TCR that had been experimentally validated to recognize the YF epitope (Liu et al. 2025), we enumerated all possible 9-mer peptides derived from the YF vaccine strain and predicted their binding affinity to HLA-A*02:01 using MixMHCpred (Tadros et al. 2025). We modelled with AF3 the 251 peptides predicted as binders (%rank < 2) in complex with the initial TCR. The cognate epitope ranked fourth among all 9-mers based on the predicted iptm_pair_mean score (Fig. 2D, Table S4C), showing how AF3 could be used to prioritize cognate epitopes of a TCR of interest out of a list of candidates.

Finally, we tested whether AF3 can predict how amino acid variants in the TCR and in the epitope impact these interactions. Using experimental single-variant scans at position 4 in the CDR3α and position 5 in the CDR3β, and at every position of the YF peptide for three YF-specific TCRs (Liu et al. 2025), we observed only a weak correlation between the experimental binding results and AF3 predictions, with many variants that abrogated TCR recognition receiving high AF3 scores (Fig. 2E&F, Table S4D&E).

Overall, our results highlight the variability of AF3 prediction accuracy for TCR-epitope recognition for different epitopes across different scenarios. In line with findings in other contexts (Feldman et al. 2026), we also observed that AF3 had limited accuracy for predicting the impact of single amino acid variants, even in the case of an epitope where AF3 is able to correctly model TCR recognition specificity.

### Antibody-specific Epitope prediction

Antibody-antigen binding and complex prediction remains an unsolved challenge in protein structural biology, largely because the antibody interface consists of unstructured loops that are not evolutionarily driven, depriving deep-learning methods of the co-evolutionary signal in multiple sequence alignments (MSAs). However, antibody development and design is highly relevant in therapeutic settings, where over 1/4 drugs that obtained FDA approval in the year 2025 were such large molecule biotherapeutics. Currently, therapeutic antibodies are expensive to develop and improved prediction models would enable better in silico screening and de novo design.

We examine how AF3 can be used to predict antibody-antigen interfaces and how the output of the predictions can be used for concrete applications such as drug development and viral escape risk assessment. We used the SARS-CoV-2 dataset from Cao et al. (Cao et al. 2022) (Fig. 3A, Table S5) due to its extensive coverage of antibodies, including their sequences, over two different SARS-CoV-2 strains - D614G (emerged Feb. 2020) and Omicron BA.1 (emerged Aug. 2021). As such, there are no Omicron receptor binding domain (RBD) structures, as obtained from the CoV-AbDab, in the AF3 training dataset (cutoff September 30th, 2021).

**Figure 3.**
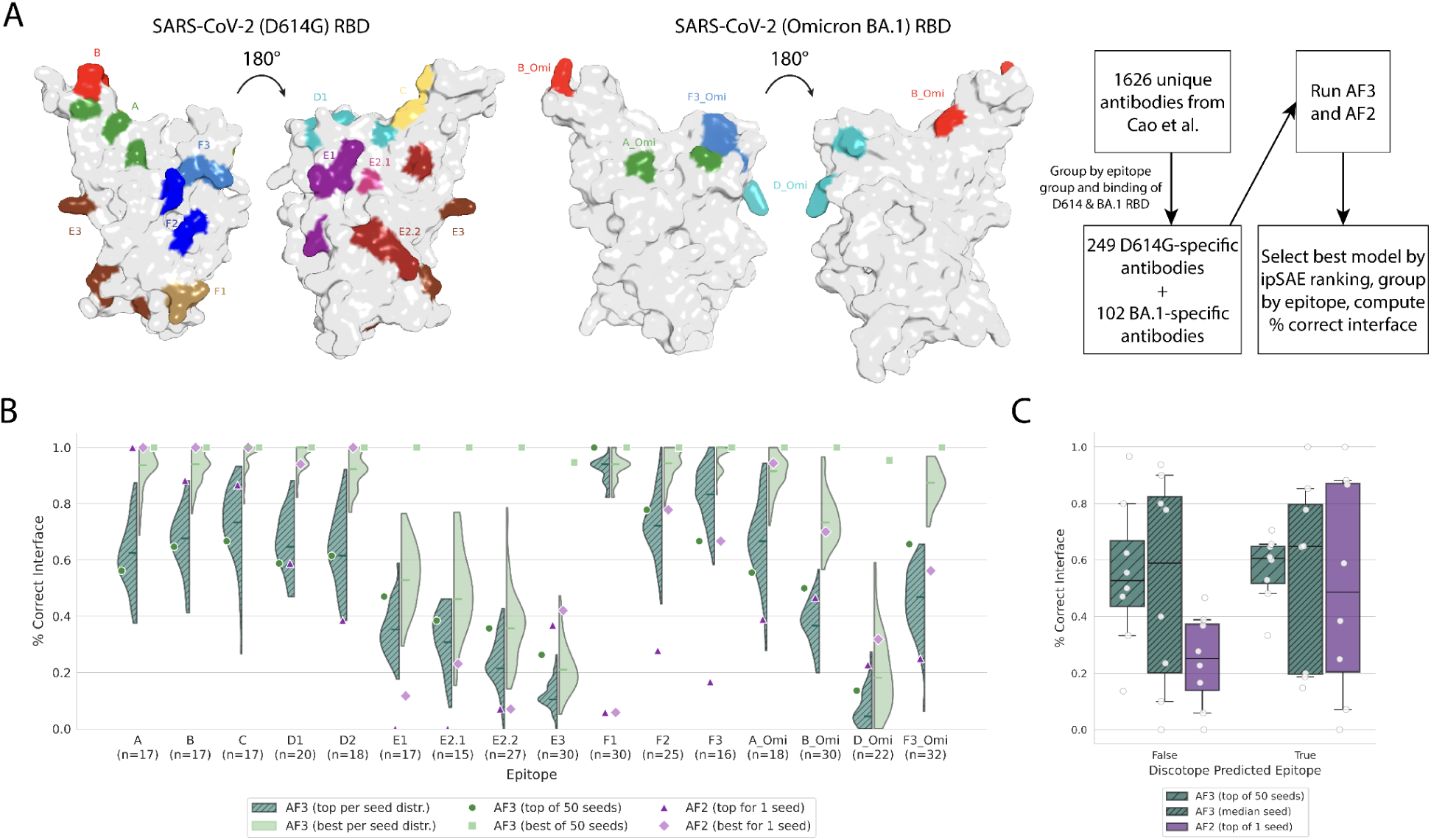
Antibody–antigen interface prediction performance on the SARS-CoV-2 RBD dataset. **A)** Epitope regions on the receptor-binding domain (RBD) of SARS-CoV-2 D614G (left) and Omicron BA.1 (right), as defined by Cao et al. **B)** Fraction of complexes with a correctly predicted interface across epitope categories. Top: models selected by each method’s own ranking (AF3 ipSAE, AF2-Multimer ranking score). Bottom: best-performing model per target, selected using ground-truth interface accuracy. **C)** Mean interface recovery for each epitope, by Discotope epitope-likesness of the true binding site, comparing AF2 and AF3 (top-ranked out of 50 seeds and the median performance across the 50 seeds).

We compared two model-selection strategies: the realistic “top” selection, ranked by each method itself, and an oracle “best” selection, using the true interface accuracy of each model (Fig. 3B). For top-ranked models, the median AF3 seed outperformed the single AF2 seed in 10 of 16 cases (Fig. 3B, upper). Notably, AF3 was relatively insensitive to the “epitope-likeness” of the true interface (measured by Discotope 3.0 (Høie et al. 2024); Fig. S5D), whereas AF2 performed better on epitope-like sites (Fig. 3C). Comparing top with best selection revealed a ranking problem. Across the 50 AF3 seeds a complex at the correct interface was almost always present (light blue circles), but AF3’s confidence and ranking rarely selected it (not the top rank). Additionally, the mean ipSAE_max over both antibody-antigen interfaces did not rescue this (Fig. 3B, lower; Fig. S5E). A similar trend is observed for AF2, however the difference is not as stark, as only one seed was run for AF2. Therefore, for some epitopes, running AF3 with 50 seeds can produce an accurate interface, but the current ranking generally fails to identify it.

Finally, but importantly, the interface recovery does not correlate with antibody binding affinity, with near equal success rates for both high and low-affinity binders (Fig. S5F). This is in line with training mode of such co-folding models, where no negative (non-binding) data is presented during training, and so the model is trained to learn to predict the correct structure and binding site given the fact that the inputs interact (Bio 2026). AF3 additionally seems to generally perform better at predicting interfaces seen in its training dataset (Fig. S5BC). Point mutations that break binding can also not be predicted with AF3 (Fig. S5G, Table S6), when evaluating on a subset of 10 antibodies each tested for binding to 75-188 RBDs differing by single point mutations. This conclusion was also proven true for similar co-folding methods such as Boltz-2 (Bio 2026).

In summary, on the Cao et al. dataset AF3 predicted more accurate interfaces than AF2 and was less dependent on epitope-likeness, though AF2 was still better for some epitopes. Running 50 seeds was sufficient before scores plateaued. As with other applications, AF3’s main weakness was specificity with its confidence modules neither reliably selected the correct interface (the pAE-based ipSAE did not help) nor correlated with binding affinity.

### Modelling protein-RNA interactions

To investigate AF3’s capacity to predict protein-RNA interactions based on model confidence scores we established a comprehensive benchmark dataset using non-structural experimental approaches. We obtained RNA targets for RNA-binding proteins (RBPs) using CLIP-seq data from the POSTAR database (Zhao et al. 2022). To map the corresponding interaction sites on the protein side, we integrated cross-linking mass spectrometry (MS) data from the RNPxl (Kramer et al. 2014) and XRNAX (Trendel et al. 2019) datasets.

Additionally, to provide an extra source of pairs with functional relevance - specifically in the context of cancer - we performed a multi-omics association analysis utilizing the CPTAC dataset (Li et al. 2023). By analyzing genomic (copy-number variations), transcriptomic, and proteomic data from 1’072 tumor samples across 10 cancer types, we identified 343 robustly correlated RBP-RNA pairs (221 positive and 122 negative correlations) that exhibited significant (p < 0.01), coordinated directional effects across all three omics levels. This yielded an extra set of high-confidence, cancer-relevant interactions involving 54 unique RBPs and 312 RNA targets for structural evaluation (Fig. 4A).

**Figure 4.**
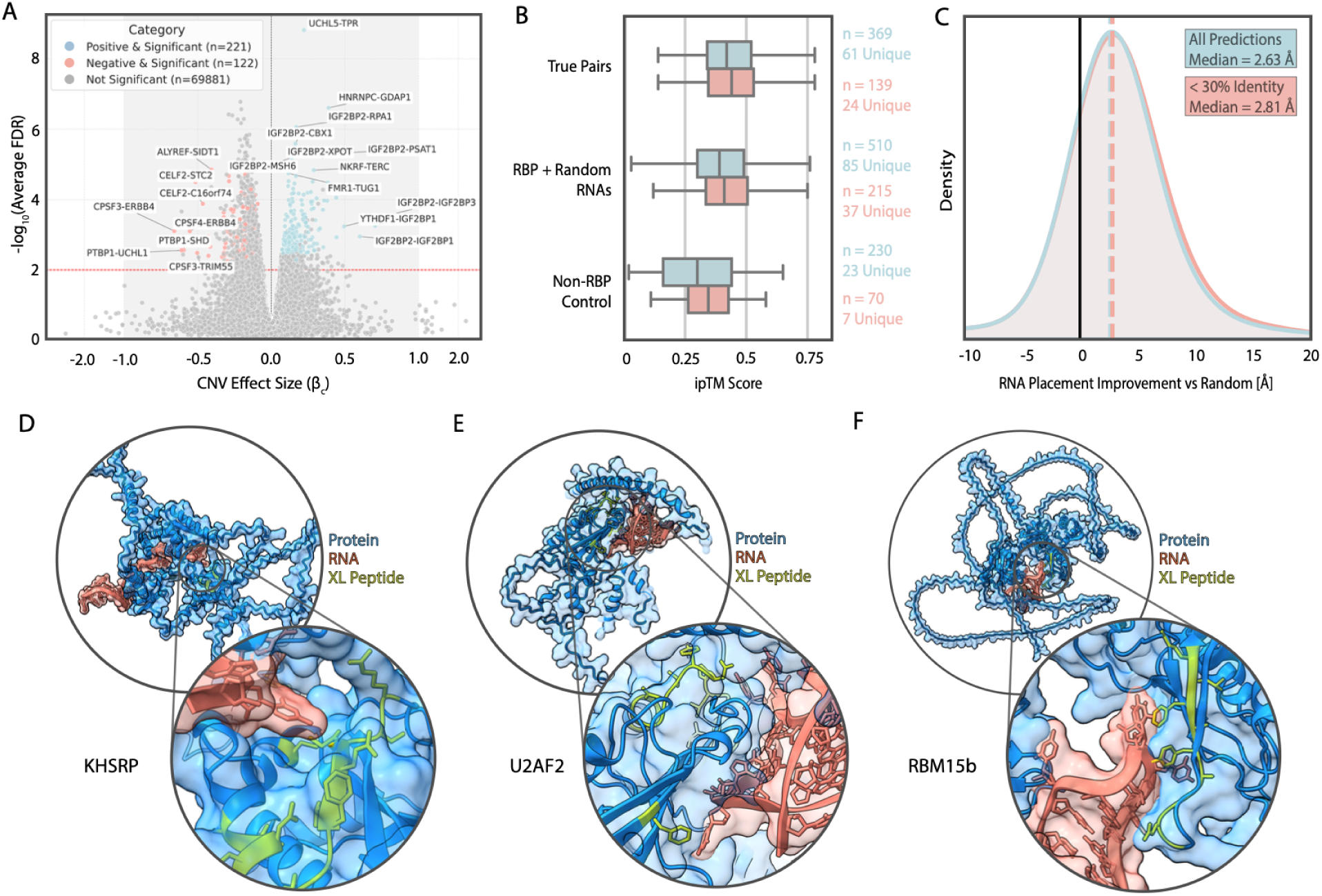
Multi-omics and structural evaluation of protein-RNA interactions. **A)** Volcano plot of the multi-omics correlation analysis. The x-axis shows the copy number variation (CNV) effect size (βc), and the y-axis displays the -log10(Average FDR). Data points represent individual RBP-RNA pairs color-coded by significance and correlation direction. **B)** Boxplot displaying AlphaFold 3 interface predicted template modeling (ipTM) scores (x-axis) evaluated across three target groups (y-axis). The data for each group is split into the full dataset and a “Low/No Homology” subset. **C)** Density plot illustrating spatial RNA placement improvement. The x-axis represents the improvement in Ångströms compared to a random baseline, and the y-axis shows the density distribution for both the full dataset and the low-homology subset. **D–F)** 3D structural representations of AlphaFold 3 predictions for KHSRP **D)**, RBM15b **E)**, and U2AF2 **F)**. Each panel visualizes the predicted protein (blue), the modeled RNA chain (red), and the highlighted experimentally-identified cross-linked peptide (XL Peptide, yellow).

To assess AF3 performance, we divided our predictions into three main groups: (1) “True Positive Pairs” consisting of validated RBPs and their experimentally or correlationally confirmed RNA targets (61 unique proteins, n = 369 predictions), (2) “RBP + Random RNA” serving as a specificity control (85 unique proteins, n = 510 predictions), and (3) “Non-RBP Control” consisting of proteins not previously described to have RNA-binding function paired with RNA (23 unique proteins, n = 230 predictions).

Overall, AF3 successfully distinguished true RBPs from non-RBPs (Fig. 4B, Table S7). The True Positive Pairs (Mean ipTM = 0.437) scored significantly higher than the Non-RBP Control group (Mean ipTM = 0.305; p < 0.001). Similarly, the RBP + Random RNA group (Mean ipTM = 0.398) also scored significantly higher than the Non-RBP Controls (p < 0.001). While True Positive Pairs scored statistically higher than the “RBP + Random RNA” group (p < 0.01), the difference is mainly driven by the additional protein diversity within the randomly paired group. Both of the groups having RNA binding proteins had higher scores than the non-RBP group. This indicates that AF3 lacks the specificity required to distinguish genuine RNA targets from random RNA sequences. Seemingly, the presence of an RNA-binding fold heavily drives the ipTM score, regardless of the exact RNA sequence provided.

We investigated whether AF3’s performance was artificially influenced by the presence of homologous structural templates in its training data using a 30% global sequence identity cut-off to define a “Low/No Homology” subset. When comparing the full dataset to the low-homology subset, we observed no significant drop in performance when templates were unavailable (Fig. 4B). In fact, the Mean ipTM slightly increased for the True Positive Pairs (0.437 vs 0.447, not significant). Surprisingly, the RBP + Random RNA (0.398 vs 0.428, p < 0.05) and the Non-RBP Control (0.305 vs 0.35, p < 0.05) group showed a significant increase in confidence scores in the absence of templates. Overall, this supports the hypothesis that AF3’s predictive ability is driven by learned representations of RNA-binding folds rather than template copying.

Despite the lack of strict RNA sequence specificity, AF3 demonstrated high accuracy in predicting the physical location of RNA binding interfaces (Fig. 4C). We evaluated 655 predictions containing 134 specific cross-link sites identified via MS (RNPxl and XRNAX, Table S8). By measuring the shortest distance between the heavy atoms of the predicted RNA chain and the experimentally validated cross-linked peptide residues, we found that AF3 placed the RNA within proximity thresholds of 3 Å, 5 Å, and 10 Å in 77%, 88%, and 91% of individual predictions, respectively. When aggregated to the protein level, this corresponds to success rates of 82%, 83%, and 87%.

Overall, AF3’s RNA placement represented a substantial improvement over random residue baselines, achieving a median placement improvement of 2.63 Å for all predictions and 2.81 Å for the low-homology subset (Fig. 4C). We visually confirmed this spatial placement in a subset of predicted 3D structures. For instance, in the structural models of the RNA-binding proteins KHSRP, RBM15b, and U2AF2, AF3 accurately models the RNA chain within the strict 3 Å threshold of the experimentally identified MS cross-link peptides (Fig. 4D-F).

### Benchmarking protein-lipid co-folding predictions

To evaluate AF3 ability to model protein-lipid co-folding, we first assembled a structurally diverse set of 12 lipid transport proteins (LTPs), covering multiple folds and lipid classes (Srinivasan et al. 2024), that were experimentally resolved with their cognate lipids. Then, we generated AF3 predictions for each protein with its cognate lipid (true positives), with a chemically diverse panel of additional lipids, and with n-alkanes (DB1 dataset, see SI). For cognate lipid binding, consistent with these complexes being in the training set, AF3 identified the correct binding cavity and reproduced the experimental binding poses with a low mean pocket-aligned lipid RMSD per atom (0.08 Å). This establishes an upper bound for AF3 performance under favourable conditions.

We computed three well-established AF3-derived metrics to assess the confidence of the co-folding predictions: min-iPAE, ipTM, and avg-pLDDT (see Methods). Across DB1, AF3 showed a strong ability to prioritize cognate lipids over other tested lipids. The co-crystallised lipid ranked first by ipTM in 6 of 12 proteins and within the top three in 11 (Fig. 5A), with comparable results for min-iPAE (7/12 first; 10/12 top three; Fig. S6). The three metrics were strongly correlated (|Pearson| ≈ 0.8; Fig. S7), with avg-pLDDT being slightly less discriminative (6/12 first; 8/12 top three; Fig. S8). We therefore selected ipTM as the primary metric for subsequent analyses given its normalized scale and straightforward interpretability. We noted that, although all true positives had ipTM > 0.8, so did many alternative lipid-protein pairs. As the specificity of most of these proteins for their co-crystallised lipids is not established, we cannot tell whether this reflects a genuine lipid-binding promiscuity or an AF3 inability to resolve lipid-binding specificity.

**Figure 5.**
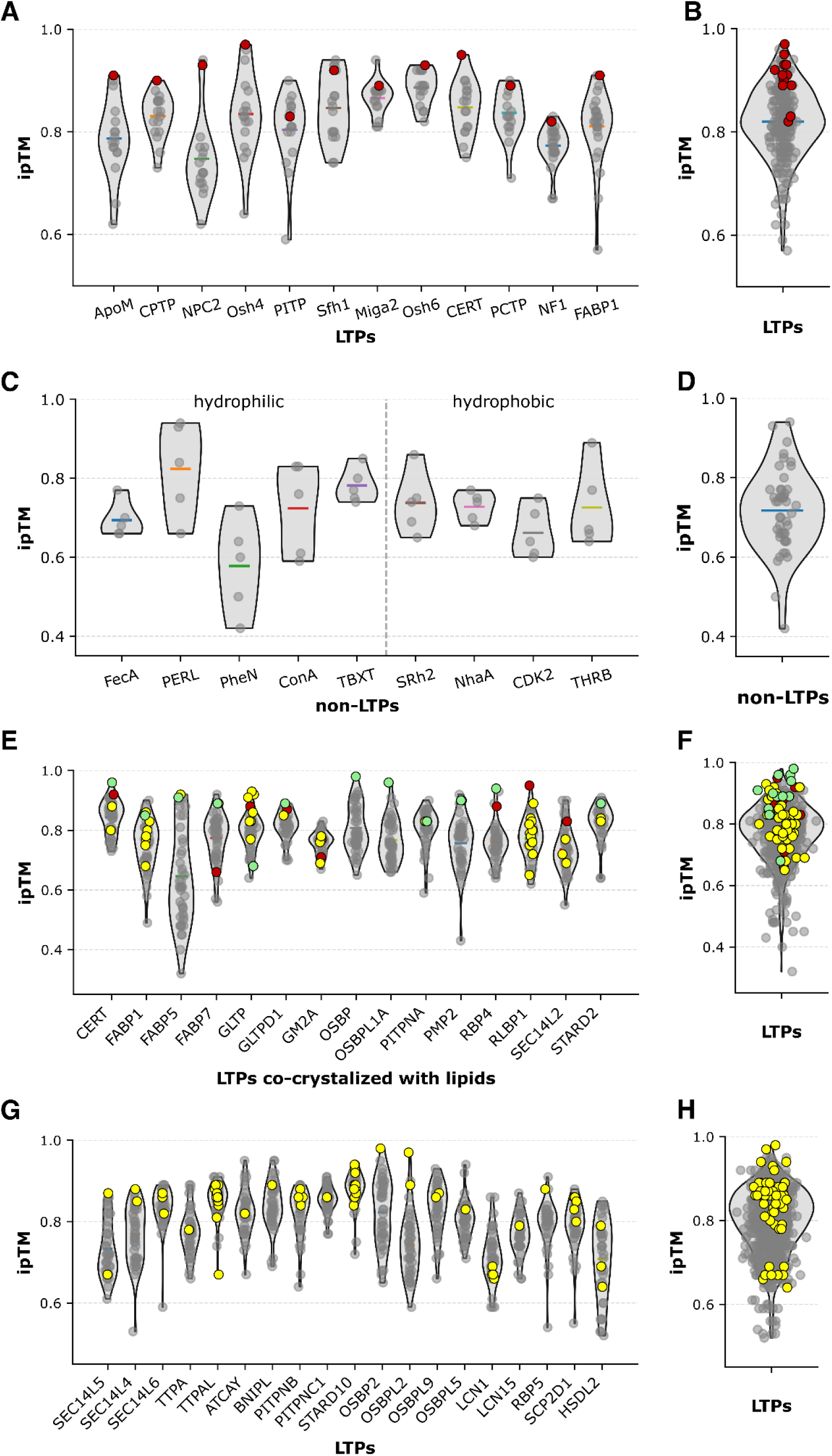
Distribution of interface predicted TM-score (ipTM) values obtained from AF3 protein-lipid co-folding predictions across multiple benchmarking datasets. Panels **A)** and **B)** correspond to DB1, **C)** and **D)** to DB2, **E)** and **F)** to DB4, and **G)** and **H)** to DB5. For each dataset, per-protein ipTM distributions are shown in the left panels **(A, C, E, and G)**, while aggregated distributions across all proteins are shown in the right panels **(B, D, F, and H)**. Across all panels, violin plots represent the distribution of ipTM scores for all predicted protein-lipid pairs. Individual data points corresponding to experimentally validated interactions are overlaid as colored markers: red dots indicate co-crystallized protein-lipid complexes (true positives derived from x-ray structures), yellow dots indicate interactions supported by mass spectrometry or high-performance thin-layer chromatography, and green dots denote interactions validated by both structural and biochemical evidence. All remaining predictions correspond to protein-lipid pairs without experimental support (putative negatives or uncharacterized interactions). The relative positioning of experimentally validated interactions within the ipTM distributions illustrates the discriminative performance of AF3 across datasets.

To more directly evaluate specificity, we tested whether AF3 can distinguish bona fide LTPs from other proteins that, although possessing large internal cavities, are unable to bind or transport lipids. To this end, we compiled a set of nine such proteins, containing both hydrophobic and hydrophilic cavities, from the literature (Chwastyk et al. 2020), created a new dataset (DB2) that included each of these proteins in complex with lipids representative of distinct families, and used AF3 to generate co-folding predictions for each of these protein-lipid pairs (50 predictions in total) (see SI).

Despite all examples being true negatives, approximately 7% of predictions yielded ipTM > 0.8, indicating a non-negligible false positive rate when large cavities are present (Fig. 5C and 5D). Combining the 12 LTP-cognate-lipid true positives with these 50 negatives (DB3), AF3 achieved an area under the precision-recall curve (AUPRC) of 0.701 (Fig. S9A). This value indicates a moderate to good discriminative power of AF3 in this case, substantially above the random baseline (which would approximate the fraction of true positives), but far from perfect classification. This implies that additional filtering or orthogonal validation would be required to unequivocally identify non-LTPs in this relatively simple scenario.

To extend the number of true positives beyond structurally resolved lipid-protein complexes, and therefore enable more robust benchmarking, we leveraged a large-scale dataset comprising 1,394 lipid-protein pairs (125 positives and 1,269 negatives), which were mainly determined by mass spectrometry (Titeca et al. 2026). This dataset was partitioned into two subsets: DB4 (615 pairs; 65 positives, 550 negatives) contained proteins with lipid-bound X-ray structures overlapping the AF3 training set, while DB5 (779 pairs; 60 positives, 719 negatives) contained proteins with no lipid-bound complexes seen during training, enabling an out-of-sample test.

Predictive performance decreased substantially in both DB4 and DB5 (Fig. 5E-H). Many experimentally validated interactions had an ipTM score below 0.8 and numerous true negatives exceeded this threshold (0.8). For DB4, AF3 achieved an AUPRC of 0.309 (Fig. S9B), with normalized enrichment factors (NEF) of 0.86 (1%), 0.35 (5%), and 0.34 (10%). For DB5, predictive performance further declined, with an AUPRC of 0.189 (Fig. S9C) and NEF values of 0.50, 0.15, and 0.18, respectively.

Three key trends emerge from this analysis. First, AF3 retains early enrichment capability (strong early enrichment in DB4 and notably weaker in DB5) indicating that a large fraction of the top-ranked protein-lipid co-folding predictions, mostly in DB4, are enriched in true positives. Second, overall precision remains limited, particularly in the fully out-of-sample setting (DB5), reflecting challenges in generalizing protein-lipid co-folding recognition beyond training distributions. Third, the performance gap between DB4 and DB5 further suggests that exposure to protein-lipid pairs seen during training influences AF3 predictive accuracy, which seems to be partially driven by the structural patterns learned rather than solely by generalizable physicochemical principles.

Overall, AF3 can very accurately reconstruct known protein-lipid interactions and identify plausible lipid binding modes, but exhibits limited specificity and generalization when applied to large-scale screening tasks. The relatively high NEF values suggest that AF3 is well suited for prioritization workflows, where a small subset of predictions can be subsequently selected for downstream validation. However, the modest AUPRC values underscore the need for complementary approaches to reduce false positives, particularly when working with chemically diverse lipid spaces and proteins for which no lipid-bound complexes were seen during AF3 training. The protein-lipid benchmark datasets are provided in Table S9.

## Discussion

We have assessed AF3 across different application areas that make use of the all-atom capabilities or reported general improvements in accuracy over AF2. AF3 extends structure prediction to biomolecules (e.g. RNA and lipids) or capabilities (e.g. chained protein covalent bounds), that are not possible in AF2 and in a few settings it performs well even where no close structural template exists. At the same time, our results show a recurring limitation in specificity, and in generalisation beyond the training data. Compared with a previous assessment of AF2 applications (Akdel et al. 2022), the overall outcome is more nuanced, suggesting a more limited application of AF3 beyond modelling of structures of monomer protein and protein-protein interactions.

Several of our results nonetheless show valuable new capability beyond simple memorization. The confidence metrics appear to enrich for well modelled ubiquitination and the loss of accuracy after removing the covalent bonds argue against simple memorisation. In the protein-RNA setting, AF3 tends to place the RNA in the correct interface even in the low-homology subset suggesting that RNA binding sites are well predicted. These results are encouraging, suggesting that AF3 can drive novel discoveries in these different areas.

The TCR-epitope results are especially encouraging given the low affinity and loop flexibility that make these interactions hard to model. Although AF3 cannot recover all TCRs binding a given epitope, epitope-specific TCRs are often ranked within the top 2% of predictions, which is typically within the number of TCRs that can be experimentally tested. All our negatives came from the same samples as the positives and were all processed in the same batches, which minimizes the risk that the observed predictive power would result from some biases in the data. In some cases it did so for epitopes lacking any homologous TCR-bound structure, suggesting such TCRs could be identified directly from repertoire sequencing of blood or tumour-infiltrating T cells. Second, we observed that AF3 could accurately prioritize the cognate epitope recognized by a specific TCRs out of all potential peptides expressed in the Yellow Fever genome. This suggests that AF3 can be used to find the cognate epitope, when the list of potential epitopes can be accurately established (Gfeller et al. 2026). Finally, we observed that AF3 had difficulties in correctly predicting how single amino acid TCR or epitope variants impact TCR-epitope recognition. This is consistent with previous findings that AF3 predictions of the impact of point mutations are not always accurate (Feldman et al. 2026).

For antibody-antigen complexes, AF3 improved interface recovery over AF2 and was less sensitive to epitope-likeness. As with TCR-epitope modeling, AF3 did not reliably predict the effect of point mutations on binding, and its confidence and ranking scores neither selected the correct interface from among 50 seeds nor correlated with binding affinity. These results show that sampling may result in models with accurate epitopes, but current confidence metrics are insufficient for correct selection.

Regarding protein-lipid co-folding, consistent with previous results reported in a recent study that assessed the structural accuracy of AF3 predicted lipid poses on a dataset of 311 high-quality lipid-protein crystal structures (Yang et al. 2026), we found that AF3 accurately reconstructs known protein-lipid complexes but displays a reduced out-of-distribution generalization. We also found that, although AF3 can identify plausible lipid binding modes, it lacks specificity and exhibits limited ability to prioritize biologically relevant protein-lipid interactions. This suggests that, in this scenario, AF3 should be mostly interpreted as a hypothesis-generating tool that must be complemented with independent computational or experimental validation.

Two recurrent themes stand out from our study: 1) AF3 often showed a lack of specificity for the prediction of biomolecular interactions; and 2) we find a high dependence of performance on training-set overlap for several applications. In the protein-RNA predictions the AF3 ipTM score did not discriminate between genuine and decoy RBP-RNA pairs. The results appear to be somewhat more specific for protein-lipid interaction predictions where ipTM scores showed a significant discrimination between true positives and decoys hits. However, for protein-lipid interactions there is a clear influence of the training data. This is consistent with studies showing a similar training-data dependence on the prediction of protein-ligand interactions (Škrinjar et al. 2025).

The practical implication of these results is that the application of AF3 and its confidence scores need to be considered differentially across different types of models. For the application areas described here, the benchmarks and accuracy estimate provided can serve as a guiding point for setting appropriate cut-offs and judging where discrimination is sufficient for routine use. For other specific applications, end-users are encouraged to design appropriate benchmark datasets for further calibration of thresholds and potential development of other confidence metrics. Such evaluations should include negative controls and need to account for training-set overlap.

This study has several limitations that are worth noting. As a collection of examples from different application areas, the benchmarking strategies are not always uniform. Some applications have benchmarks that rely on indirect evidence such as cross-linking data or large-scale experimental screening methods, while others rely on direct structural models.

While we tried to consistently probe for training-set memorization, this was not always done in the same way. We also performed our analysis only on AF3 and did not attempt to compare with open reproductions AF3. We note that the benchmarks gathered for this work should be useful for the community for testing of future improvements of these methods.

In summary, AlphaFold3 advances structure prediction to a wider range of application areas and it can perform well in different settings. The value of AF3, much like other computational models, will be greatest when used as one component within a broader workflow that includes appropriate controls and experimental validation.

## Acknowledgments

SV acknowledges support by the Swiss National Science Foundation (grant CR00I5-236020). YL and DG acknowledge support by the Swiss National Science Foundation (grant 320030-231333). PB is supported by the Helmut Horten Stiftung and the ETH Zurich Foundation.

## Author Contributions

YL and JR performed and analyzed the AF3 predictions for TCR-epitope pairs. JR implemented the AF3 pipeline locally. YL compiled the data and generated the Figure. DG designed and supervised the analysis of TCR-epitope predictions. For protein-RNA, LL ran and studied the structural predictions and wrote the text, PB supervised the analysis. For modelling of ubiquitinated proteins, EK contributed towards the curation of the benchmark, RH and JvG performed and analyzed the structural modelling with assistance from JJ, JvG made figures and wrote the text, and PB supervised the work. For antibody-epitome modelling OC performed and analyzed the structural modelling, wrote the text and JD supervised the work. For protein-lipid co-folding, PC and DA performed the AF3 predictions, PC analyzed the AF3 predictions and generated the figures, PC and SV designed the study and the analyses and wrote the text. PB coordinated the collaborative project with the help of SV, JD, and DG. All authors contributed with revisions to the manuscript.

## Methods

### Modeling structures of ubiquitinated proteins in AF3

#### Computational analysis

All analyses were performed using Python (v3.13.2) and R (v4.5.1) within Visual Studio Code (v1.107.1). Data processing was conducted primarily in Python unless otherwise specified. Plots were generated in R using ggplot2 (v3.5.2). Protein structures were visualised in PyMOL (v3.1.4). Unless otherwise specified, all code used to analyse data is accessible at https://github.com/JulianvanGerwen/ubnAF3Benchmark.

#### Predicting ubiquitinated protein structures in AlphaFold3

AlphaFold3 v3.0.1 was used with the default settings, except for the ubiquitin bond. We treat the C-terminal glycine (Gly76) on the ubiquitin as a ligand, and explicitly specify covalent bonds to the lysine on the target protein, and to the (truncated) ubiquitin chain using bondedAtomPairs. Custom scripts were written to automate this, available at https://github.com/jurgjn/af3-polymer-bonds.

#### Construction and structural prediction of the benchmarking dataset

All Protein Data Bank (PDB) entries containing a ubiquitin-like domain (InterPro accession: IPR019956), encompassing ubiquitin and ubiquitin-like proteins, were extracted through an API call. For each entry, the first biological assembly was downloaded as a CIF file.

Covalent linkages between ubiquitin (or ubiquitin-like proteins) and partner proteins were identified using a combination of geometric searches and manual curation. First, a stringent search was performed to detect canonical isopeptide bond chemistry by identifying atom pairs in which the carboxyl carbon of a glycine residue in ubiquitin and the epsilon nitrogen of a lysine residue in a separate protein chain were separated by less than 2 Å. Next, a relaxed search was conducted to capture non-canonical bond chemistries by identifying cases in which any atom of the C-terminal residue of ubiquitin was located within 2.5 Å of any atom in another protein chain. This relaxed search was extended to include structures in which a small-molecule ligand covalently bridges two proteins, which is often the case when non-canonical amino acids are used to introduce ubiquitin-protein bonds. Finally, additional covalent linkages were identified through manual inspection of structures containing a ubiquitin-like domain.

Identified linkages were filtered to retain chemically plausible bond types, specifically Gly–Lys carbon–nitrogen linkages, bonds involving a cysteine thiol, glycine carbon to serine hydroxyl linkages, glycine carbon to non-canonical amino acids, and linkages between two non-canonical amino acids.

Protein and DNA sequences were extracted directly from the experimental CIF files. All non-proteinogenic amino acids were converted to the most similar proteinogenic amino acids prior to modelling. For example, phosphoserine (SEP) was converted to serine (SER), and L-thialysine (SLZ) was converted to lysine (LYS). Structures with more than 5,1000 residues were excluded as these cannot be modelled in AlphaFold3. Structures were then predicted with covalent bonds included as described above.

#### Comparing predicted and experimental structures

For each experimental structure, five AlphaFold3 models were generated and compared to the corresponding experimental structure. First, DockQ (version 2.1.3) was run on each pair of experimental and predicted structures, automatically identifying the best chain correspondence when multiple homologous chains are present. Next, using protein–protein interfaces detected by DockQ, structures were filtered to exclude cases in which ubiquitin did not form a substantial interface with any other protein, defined as an interface size of ≤ 10 atoms. Such cases likely represent structures in which ubiquitin is flexibly attached and are less meaningful to predict with AlphaFold3. Several structures were also manually removed because extraction of protein-protein covalent bonds or protein sequences from PDB files was unsuccessful, or because DockQ failed to identify the correct chain correspondence. Finally, predicted and experimental structures were aligned in PyMOL (version 3.1.4) using the align function with cycles = 0 and the optimal chain ordering determined by DockQ, yielding root-mean-square deviation (RMSD) values. Alignment was performed either across the entire structure or for each pair of covalently bonded proteins. In structures containing multiple bonded protein pairs, the mean RMSD across all pairs was calculated.

#### Searching for structures dissimilar to AlphaFold3 training data

We filtered our benchmark dataset for structures where the bonded protein-ubiquitin pair was not similar to any bonded protein pair present in a structure released before the cutoff date for AlphaFold3 training. First, we extracted pairs of ubiquitin (or a ubiquitin-like protein) covalently bound to another protein from our benchmark dataset, filtering for structures released after the AlphaFold3 training cutoff date (30th September 2021). For each sequence in the protein pair, we then searched for similar sequences in experimental structures from the PDB. Briefly, for each input protein sequence, a query FASTA file was generated and searched against UniRef90 using jackhmmer with one iteration and per-job parallelization. The resulting multiple sequence alignment was then used by hmmbuild to construct a profile HMM. This HMM was subsequently searched against the PDB sequence database with hmmsearch using permissive AlphaFold3-like thresholds, including relaxed filter settings (--F1 0.1 --F2 0.1 --F3 0.1) and generous cutoffs (-E 100 --domE 100 --incE 100 --incdomE 100) to retain a broad set of potential hits. We then filtered the potential hits for those with a sequence identity of 30% or greater relative to the input sequence. For each ubiquitin-protein pair, this filtering recovered sequences in the PDB similar to either ubiquitin or the covalently bound protein. Finally, we identified cases where a pair of similar sequences were also covalently bound to each other within our entire benchmark dataset, only considering structures released before the AlphaFold3 training cutoff. When no such case could not be found, the ubiquitin-protein pair was classified as novel compared to structures released before the AlphaFold3 training cutoff.

### Modelling protein-RNA interactions in AF3

#### Data Integration and Multi-Omics Correlation

Analysis A primary benchmark dataset was generated by integrating CLIP-seq data from POSTAR (Zhao et al. 2022) with MS-derived protein cross-link data from RNPxl (https://doi.org/10.1038/nmeth.3092) and XRNAX (Sarnowski et al. 2022). To define an extra source of interactions with high functional relevance in cancer, genomic, transcriptomic, and proteomic data from 1,072 tumor samples (across 10 cancer types) were retrieved from the CPTAC database (Li et al. 2023). Ordinary least squares regression was performed using three null and alternative model pairs to evaluate the effect of an RBP’s copy number variation (CNV), transcript, or protein level on its target’s transcript abundance. The models controlled for age, sex, and cancer type. RBP-RNA pairs that yielded an adjusted significance of p < 0.01 across all three omics levels with matching effect directions were defined as robustly correlated functional pairs.

#### AlphaFold 3 Structural Prediction and Classification

Complexes were predicted using an installation of AlphaFold 3 on the Euler high-performance computing cluster. Prediction targets were structured into three main groups to evaluate model specificity and accuracy. The True Positive Pairs group consisted of experimentally validated interactions, combining pairs with highly confident MS cross-links and matched CLIP RNA alongside the functionally correlated CPTAC subset, resulting in 369 structural predictions across 61 unique RBPs. To establish a specificity control, the RBP + Random RNA group was generated by taking 69 unique RBPs (originally 70, one duplicate removed) and predicting each against 5 randomly assigned RNA sequences, yielding 345 total predictions. Additionally, the 16 RBPs from the crosslinking dataset were paired with 10 RNAs that came from other crosslinked RBPs which resulted in the final 85 unique proteins and 510 predictions of this category. Finally, the Non-RBP Controls group initially utilized 25 proteins manually verified to lack RNA-binding annotations. Each was paired with 10 random RNA sequences to generate 250 predictions. After removing failed predictions, 23 unique proteins (230 predictions) remained. RNA lengths were standardized to 25 nucleotides across all groups to allow testing of sequence and structural specificity without length-induced bias. Model confidence was assessed utilizing the interface predicted template modeling (ipTM) score and overall Ranking Score. Because Shapiro-Wilk and Levene’s tests indicated violations of normal distribution and unequal variance, statistical differences in ipTM scores among the three main groups were evaluated using the global Kruskal-Wallis test followed by Dunn’s post-hoc test with False Discovery Rate (FDR) correction for multiple comparisons.

#### Template Search Analysis

To evaluate the reliance of AF3 on homologous templates, the HMMER pipeline was utilized. Searches were performed using the reference sequence database employed by AF3 for HMM building against the PDB database, strictly restricted to structures released before the AF3 training cut-off date of September 30, 2021. A global sequence identity cut-off of 30% was applied to partition the data into “All Proteins” and “Low/No Homology” subsets for rigorous evaluation. Statistical comparisons between the “All Proteins” and “Low/No Homology” sets within each specific group were performed using the Mann-Whitney U Test.

#### Interface Placement and Distance Evaluation

The accuracy of AF3’s interface placement was validated against concatenated cross-link site data from the RNPxl and XRNAX datasets. An AF3-predicted residue was defined as a binding residue if it was located within 3 Å, 5 Å, or 10 Å of any heavy atom belonging to the RNA chain. These predicted residues were subsequently compared to the MS-identified ground truth residues to calculate placement success rates. Improvements in spatial placement were aggregated and calculated against number-matched random residue baselines.

### Benchmarking TCR-epitope interaction predictions with AF3

#### Predicting interactions with AF3

To predict TCR-epitope interactions, the full MHC, b2m, peptide, TCRα and TCRβ sequences were inputted to AF3. AF3 was run on a local server with GPU acceleration, considering a single seed. To speed up computations, pre-computed paired and unpaired MSAs were provided for the MHC, b2m, TCRα and TCRβ. For each prediction, the best-ranking model was kept, and the mean of the interface predicted template modeling (ipTM) scores between the chain pairs peptide-TCRα, peptide-TCRβ, HLA-TCRα and HLA-TCRβ was used as final confidence score.

#### Epitope-specific and baseline TCR repertoire datasets

Yellow Fever epitope-specific and baseline TCR repertoires were obtained from ref. (Liu et al. 2025). In this study, CD8⁺ T cells were isolated from a donor vaccinated against yellow fever. A fraction of the CD8⁺ T cells was stimulated in vitro using antigen-presenting cells loaded with the HLA-A*02:01-restricted peptide LLWNGPMAV. Following stimulation, antigen-specific T cells were identified and sorted using HLA-A*02:01_LLWNGPMAV tetramers. The non-stimulated CD8⁺ T cells (for baseline TCR repertoire) and the tetramer-sorted antigen-specific T cells were subjected to paired TCR sequencing using the 10x Genomics platform (10x Genomics; see (Liu et al. 2025), Table S4A).

TCR repertoires were also retrieved from the study by Gumpert and colleagues (Gumpert et al. 2026), where recognition of three neo-epitopes had been characterized. The epitope-specific T cells and epitope-negative T cells were sorted separately based on peptide-MHC multimers and their TCRs were sequenced with the 10X pipeline (Table S4B). A few TCR clones were found both in the positive and negative fractions and were kept here as epitope-specific. Four TCRs were further functionally characterized. These are referred to as TCR1-4 in the current manuscript, with the corresponding names in the original study: TCR1 = T157.3 (TRAV24, CDR3α: CALRDNYGQNFVF, TRAJ26, TRBV4-1, CDR3β: CASSQDRGFSQPQHF, TRBJ1-5), TCR2 = T157.1 (TRAV12-2, CDR3α: CAVGGYQKVTF, TRAJ13, TRBV6-1, CDR3β: CASSEKGRGRVTGELFF, TRBJ2-2), TCR3 = T157.2 (TRAV12-2, CDR3α: CAVKTFGGGNKLTF, TRAJ10, TRBV4-1, CDR3β: CASSQDIAKGGNTIYF, TRBJ1-2) and TCR4 = T112.1 (TRAV24, CDR3α: CASPTGNQFYF, TRAJ49, TRBV10-2, CDR3β: CASSDLAGHTYEQYF, TRBJ2-7).

#### Predicting the peptide recognized by a Yellow Fever specific TCR

To evaluate whether AF3 can identify the peptide recognized by a given TCR from a list of candidate peptides, we retrieved the amino acid sequence of the YF virus polyprotein (UniProt: P03314, POLG_YEFV1) and generated all possible overlapping 9-mer peptides. We then used MixMHCpred (Tadros et al. 2025) to predict peptide binding to HLA-A*02:01 and excluded peptides with predicted binding %rank > 2. The remaining candidate peptides were used for prediction with AF3 in complex with HLA-A*02:01, b2M, and the YF-TCR1 TCR (TRAV12-2, TRAJ30, CDR3α: CAVGDDKIIF; TRBV28, TRBJ2-7, CDR3β: CASTPQTAYEQYF) (Table S4C).

#### Predicting the effect on epitope recognition of single amino acid substitutions in TCRs

Binding measurements between TCR variants and the Yellow Fever epitope were retrieved from ref. (Liu et al. 2025) (Table S4D). These include all single amino acid substitutions at position P4 of CDR3α and position P5 of CDR3β of the YF-TCR1, except cysteine. Each TCR variant was modeled in complex with the LLWNGPMAV–HLA-A*02:01 complex using AF3.

#### Predicting the effect of single amino acid substitutions in epitope using AF3

X-scan data from ref. (Liu et al. 2025) were used (Table S4E). For three different TCRs specific to the HLA-A*02:01-restricted YF epitope LLWNGPMAV (YF-TCR1, YF-TCR2, YF-TCR3), all single-amino acid variants were generated at each position of the epitope and TCR recognition was experimentally assessed. Each peptide variant was modeled in complex with the three TCRs using AF3.

#### The sequence of the three TCRs are

YF-TCR1: TRAV12-2, TRAJ30, CAVGDDKIIF, TRBV28, TRBJ2-7, CASTPQTAYEQYF YF-TCR2: TRAV12-2, TRAJ30, CAVNPDKIIF, TRBV4-2, TRBJ1-4, CASSQEDRGPEKLFF YF-TCR3: TRAV12-2, TRAJ30, CAAGDDKIIF, TRBV29-1, TRBJ2-1, CSVATSGGSNEQFF

### Protein-lipid co-folding

#### Benchmark datasets

To assess the ability of AF3 to prioritize cognate lipids in in-silico protein-lipid co-folding studies and its discriminative power between bona fide LTPs and non-LTPs containing large internal cavities, we compiled diverse benchmark datasets. These datasets were created by pairing proteins having distinct folds (previously used in (Srinivasan et al. 2024) and (Chwastyk et al. 2020)) with lipids from major families and with selected n-alkanes. The latter were chosen as controls for pure nonspecific hydrophobic interactions. Known LTPs paired with their co-crystallized lipids were considered as true positives, while non-LTPs paired with lipids were labeled as true negatives.

To evaluate AF3 performance beyond protein-lipid complexes with experimentally resolved structures, we also leveraged a large-scale experimental dataset from a previous study (Titeca et al. 2026) which characterized lipid binding across human LTPs using MS and HPTLC. This experimental dataset was curated by excluding proteins with excessively large cavities/tunnels —such as those suspected to bridge membranes and form direct channels for lipid transport between organelles. This resulted in a dataset containing 1,394 experimentally supported protein-lipid interaction pairs (125 positives and 1,269 negatives) in total. This dataset was partitioned into two subsets to investigate the impact of training-set exposure on predictive performance: (i) a “training-overlap subset”, containing proteins with one or more lipid-bound structures overlapping with entries used during AF3 training, and (ii) an “out-of-distribution subset”, including all proteins for which no lipid-bound structures were represented during AF3 training.

#### AF3 protein–lipid co-folding predictions

All co-folding predictions were carried out using a locally installed version of AF3. For the prediction of each protein-lipid pair, no templates were employed, and only the protein amino acid sequence and lipid SMILES were supplied as input. Inference was performed under default parameter settings and using a single random seed from which five diffusion samples were generated. The resulting five protein-lipid co-folding models were subsequently ranked using the AF3 internally calculated ranking_score values. Only the top scoring model (highest ranking_score) was retained for downstream analysis.

#### Structural evaluation and confidence metrics for protein–lipid interaction assessment

The quality of the lipid binding poses predicted when LTPs were modeled with co-crystallized lipids was measured using a mean pocket-aligned lipid RMSD per atom. Pocket alignment between x-ray and co-folded structures was performed using all backbone atoms belonging to protein residues within a 10 Å cutoff of any of the lipid heavy atoms.

We use three AF3-derived confidence metrics to assess the reliability of the predicted complexes: (i) min-iPAE (minimum predicted aligned error between lipid and protein interface atoms) (Omidi et al. 2024), ipTM (interface predicted TM-score) (Evans et al. 2021), and avg-pLDDT (mean predicted local distance difference test computed across lipid atoms) (Mariani et al. 2013). Lower min-iPAE values and higher ipTM and avg-pLDDT values indicate greater confidence in the predicted protein–lipid interaction. Correlation between confidence metrics was quantified using Pearson correlation coefficients.

#### Predictive performance and early enrichment analysis

Two complementary metrics were used to quantitatively determine AF3 predictive accuracy. AF3 discriminative power between true and false positives was quantified using the area under the precision-recall curve (AUPRC), while its ability to place true positives between the top ranked predictions (early recognition) was measured using normalized enrichment factors (NEF) (Liu et al. 2019). We evaluated NEF at 1%, 5%, and 10% thresholds to capture performance across different screening depths.

### Antibody-antigen complex prediction

To determine how many seeds were necessary to obtain stable results, two antibodies (Abs) from the dataset, BD55-1809 and OC220225-SN0290, exclusively binding to different SARS-CoV-2 variants, D614G and Omicron BA.1 respectively, were selected and AF3 was run with 100 seeds for both antibodies in complex with their binding RBD strain, their non-binding RBD strain and a single point mutant from the dataset corresponding to the highest mutation escape score for this antibody (455D score = 0.944 and 501A score = 0.379, respectively). The individual scores for 1, 50 and 100 seeds are shown in the table (Fig. S5A), where the mean over 10 bootstraps was computed for the number of seeds <100. B=binding, NB=non-binding, Mut=single point mutant with high escape score. The difference in predicted scores between the non-binding strain or mutant and the binding strain was computed for the range of 1-100 seeds (Fig. S5A). We conclude that running 50 is sufficient for obtaining stable AF3 in silico scores, as from 50 seeds on, no improvement is observed. The subsequent analyses were therefore run with 50 seeds for each complex.

To evaluate how good AF3 is at predicting the correct binding interface, we selected a total of 249 and 102 antibodies binding to the D614G and Omicron BA.1 strain, respectively. We selected the antibodies to cover all the sources (published DOI, or plasma donor-extracted) and all of the epitope categories identified in the paper. AF3 (50 seeds, 5 samples, 250 models per target) and AF2-Multimer (1 seed, 5 models per target) were run for all of these antibodies in complex with their respective RBD strain. A complex being predicted with the correct interface is defined here as having at least one residue within +/- 2 residues (in sequence) of the true interface residues defined in Cao et al.

## Supplementary Figures and Tables

**Figure S1.**
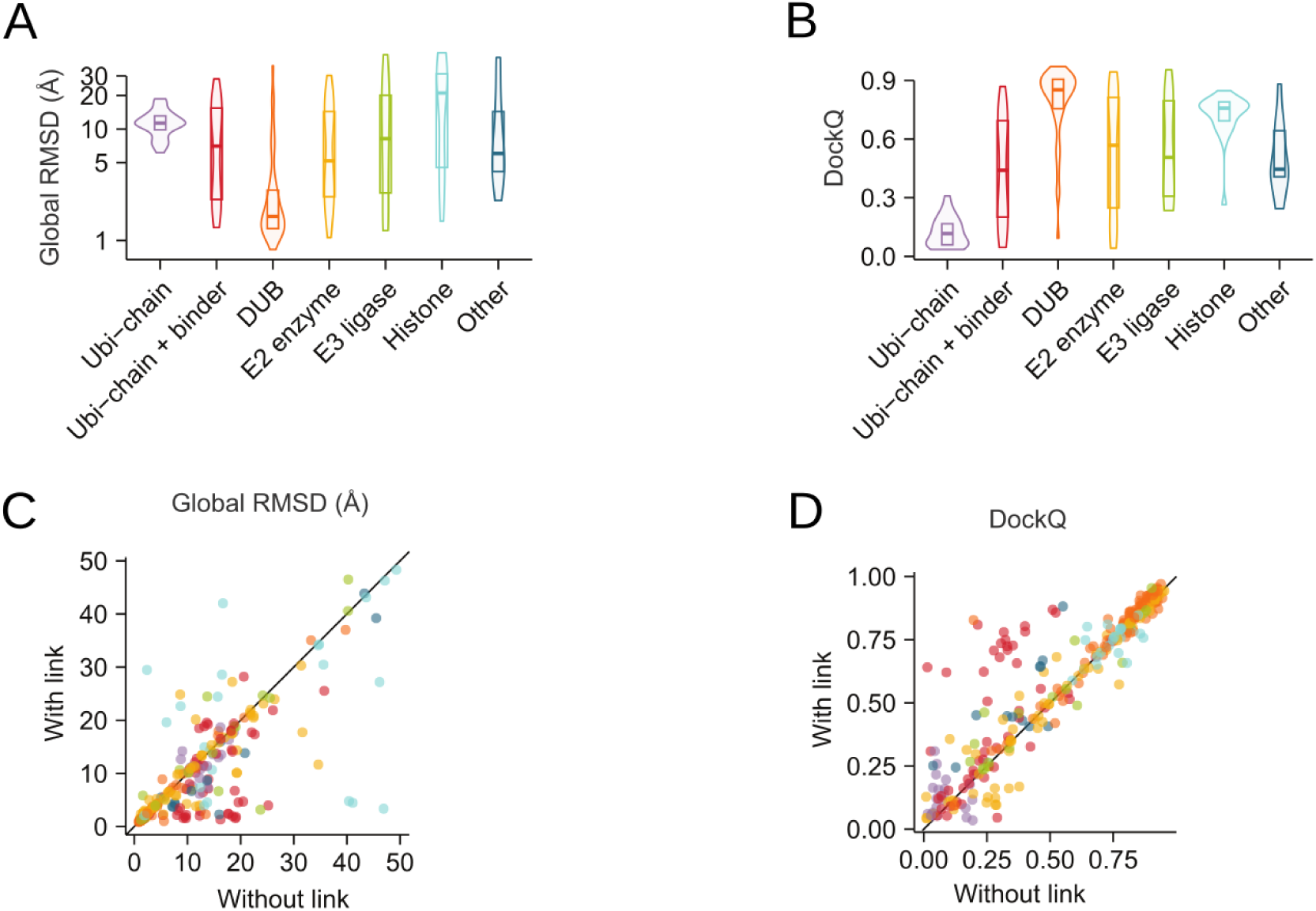
Alternative evaluation metrics for ubi-structure predictions. **A)** Root-mean-square deviation (RMSD) comparing predicted and experimental structures, calculated over all proteins. RMSD was averaged across five AlphaFold3 models (see Methods). **B)** As in **A)**, using the DockQ metric. **C-D)** Average **C)** RMSD (calculated on all proteins) and **D)** DockQ across five AlphaFold3 models with protein-ubiquitin bonds included or excluded.

**Figure S2.**
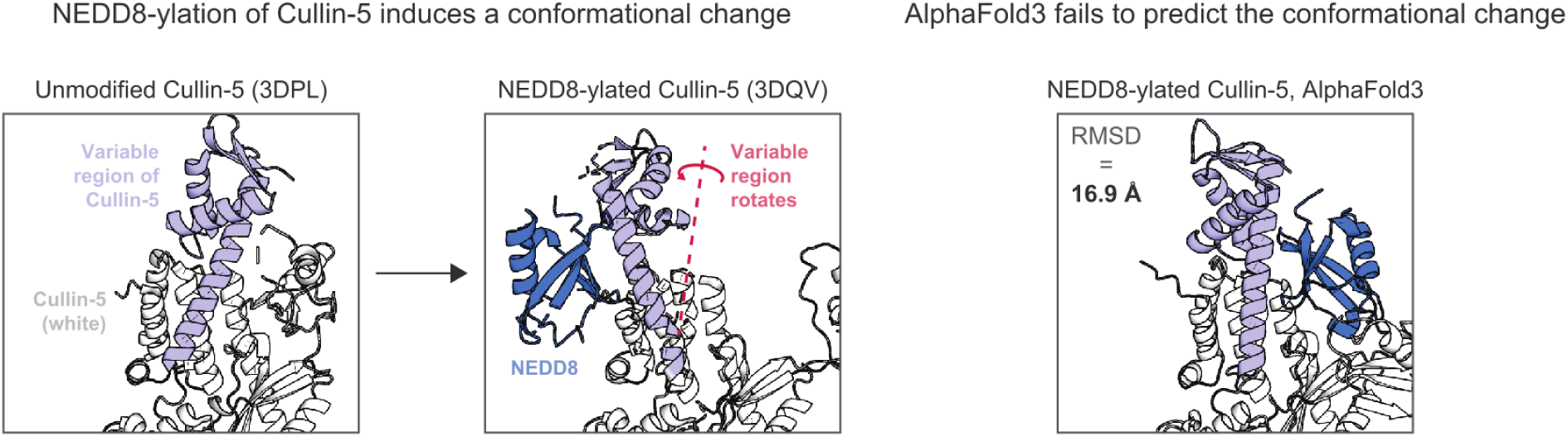
Prediction of the NEDD8-ylation-induced conformational change in Cullin-5. Left: Structures of unmodified and NEDD8-ylated Cullin-5. The region of Cullin-5 that displays a conformational challenge in response to NEDD8-ylation is coloured light blue. Right: AlphaFold3 predictions of NEDD8-ylated Cullin-5. The RMSD compared to the experimental structure (3DQV) is shown.

**Figure S3.**
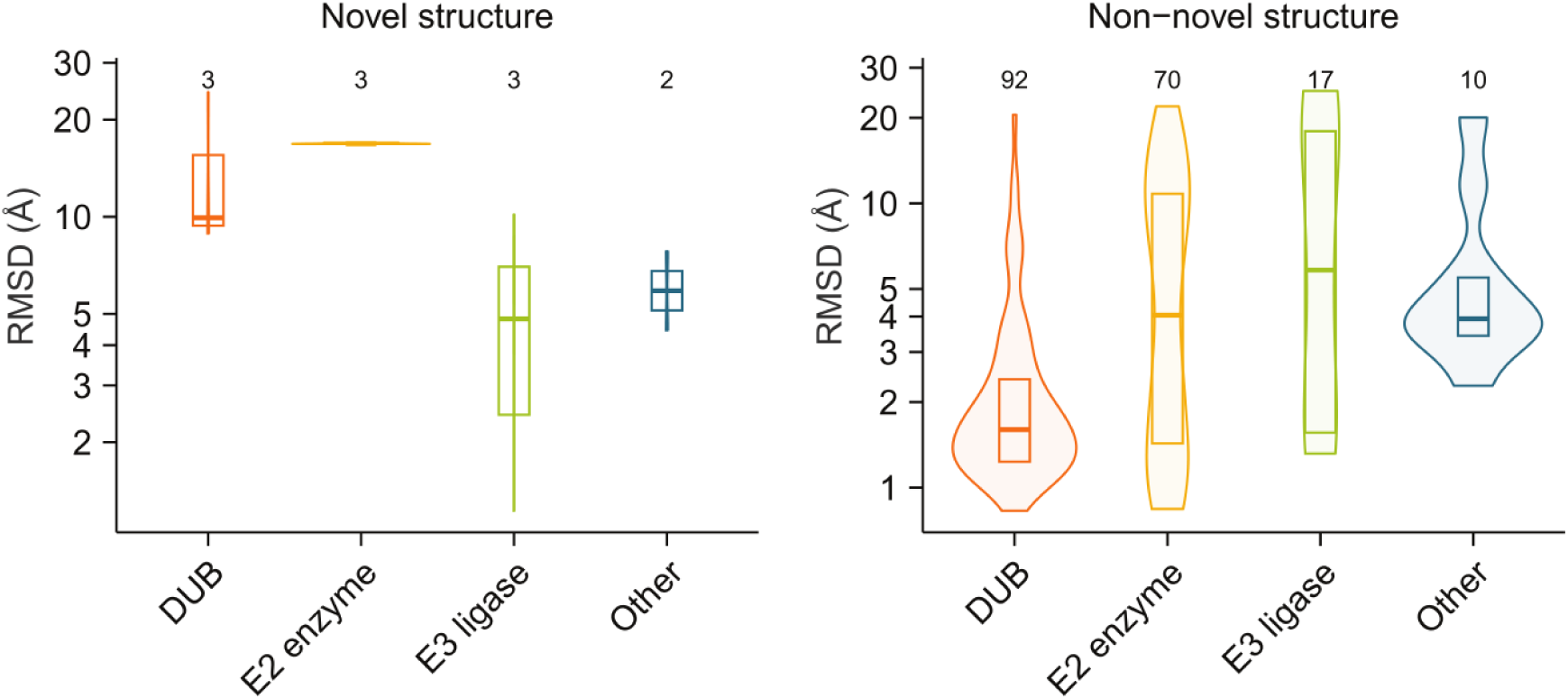
Searching for structures dissimilar to AlphaFold3 training data. The benchmark dataset was filtered for structures where the bonded protein-ubiquitin pair was not similar to any bonded protein pair present in a structure released before the cutoff date for AlphaFold3 training (see Methods). For the resulting structures (“Novel structure”) the root-mean-square deviation (RMSD) comparing predicted and experimental structures is shown. The RMSD for all remaining structures in the same structural groups (“Non-novel structure”) is also shown. RMSD was calculated on pairs of covalently bonded proteins, using the average when more than two proteins were bonded. RMSD was averaged across five AlphaFold3 models (see Methods). The RMSD values for these structures are also shown in Fig. 1C.

**Figure S4.**
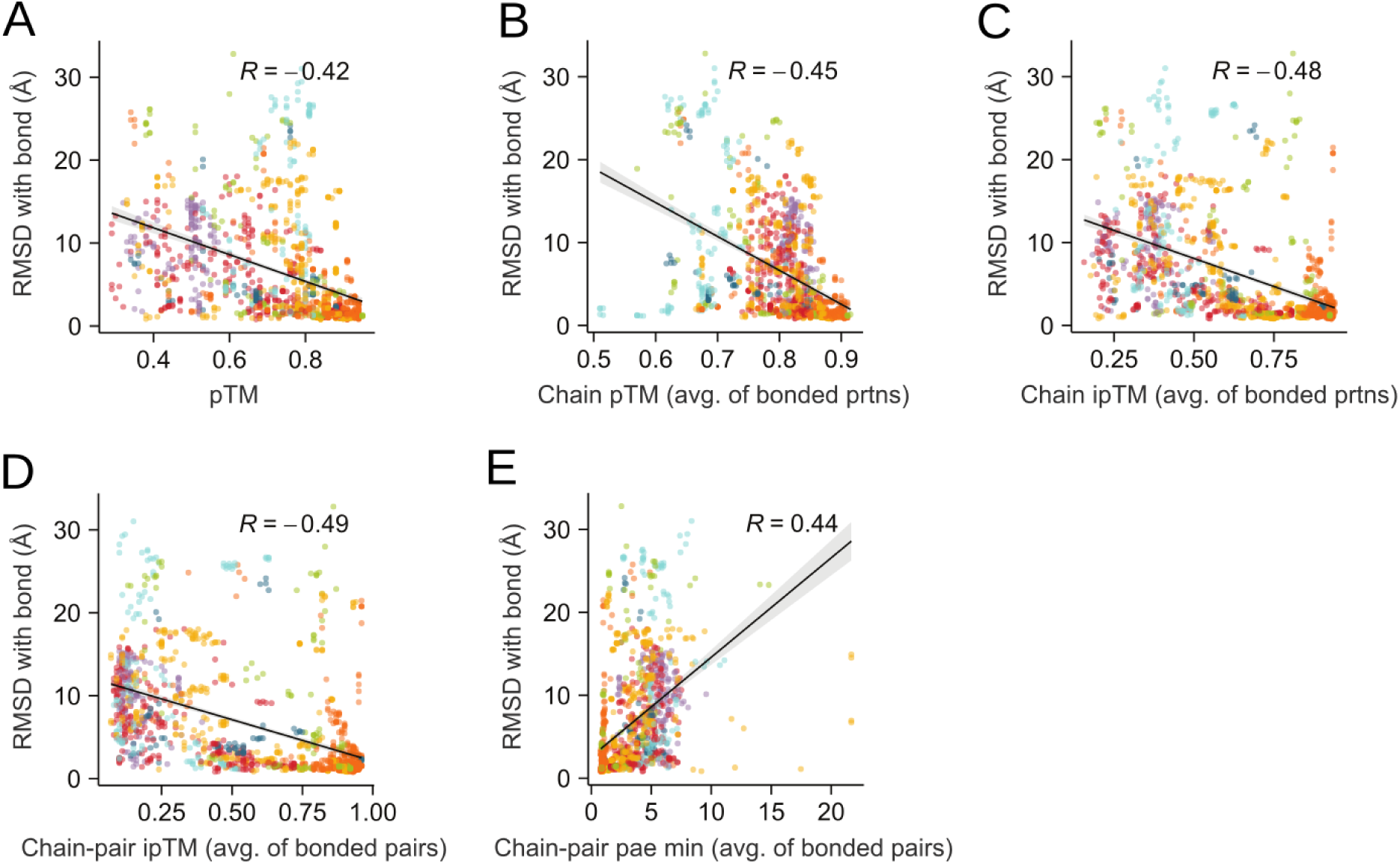
Alternative confidence metrics for ubi-structure predictions. Relationship between RMSD (calculated on pairs of bonded proteins, shown in Fig. 1h) and AlphaFold3 model confidence metrics Protein-ubiquitin bonds were included. For each structure, the five AlphaFold3 models are shown separately. Linear regression line is shown with 95% confidence interval shaded, and Pearson’s correlation coefficient is shown. **A)** Structure-level pTM; **B)** Chain-level pTM, averaged across proteins in covalent bonds; C) Chain-level ipTM, averaged across proteins in covalent bonds; D) Chain pair-level ipTM, averaged across pairs of bonded proteins; E) Chain pair-level minimum pae, averaged across pairs of bonded proteins

**Figure S5.**
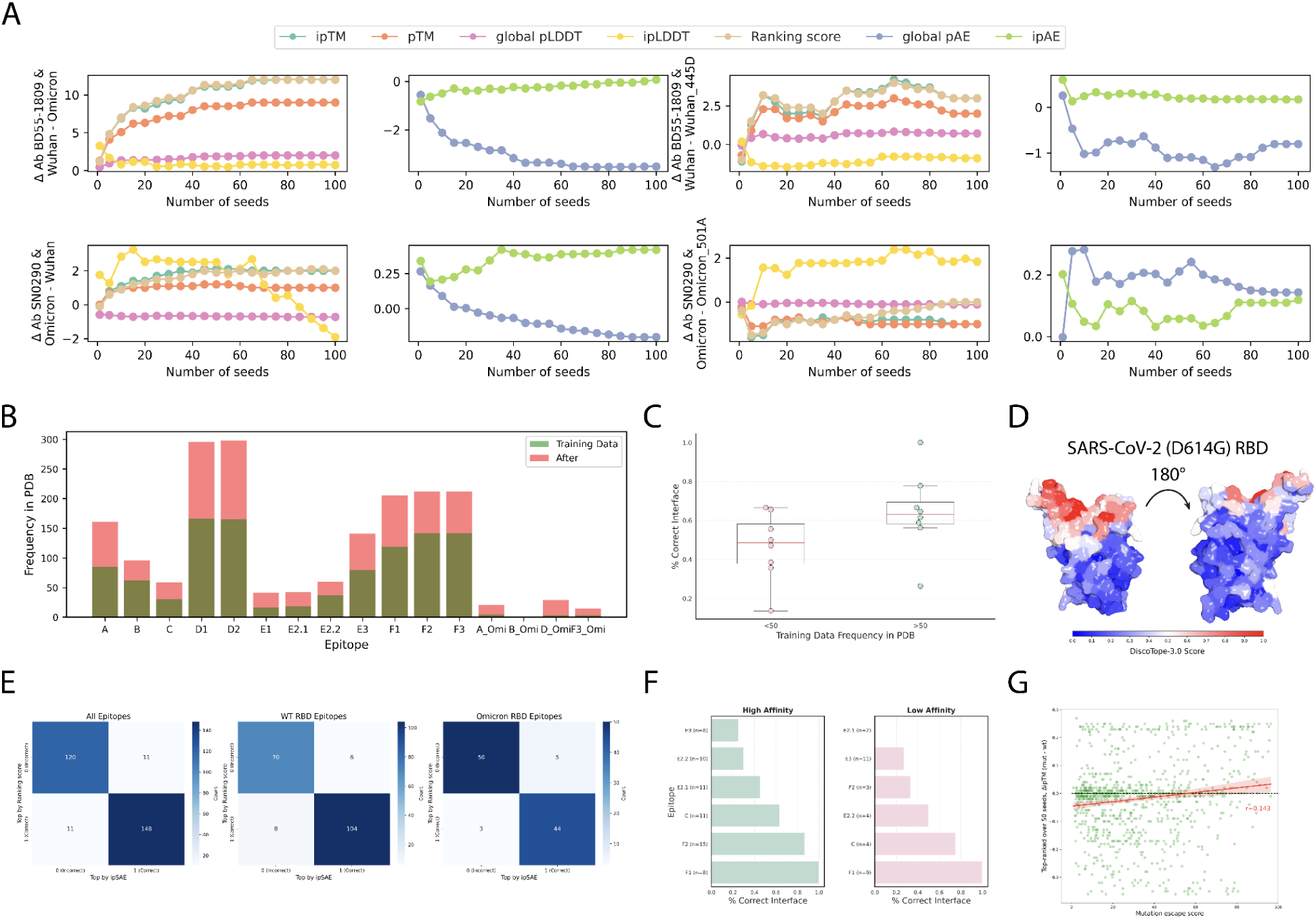
Supplementary analyses of AlphaFold3 antibody-antigen prediction. **A)** Convergence of AF3 confidence metrics with increasing seed number (1-100) for two representative antibodies: BD55-1809 (D614G-specific; top row) and OC220225-SN0290 (Omicron BA.1-specific; bottom row). For each antibody, Δ metrics were computed between the binding RBD and either the non-binding RBD (left two plots per row) or a single point mutant with the highest mutation escape score in the dataset (Wuhan RBD 445D for BD55-1809; Omicron RBD 501A for SN0290; right two plots per row). Metrics where high is better (left plots): ipTM, pTM, global pLDDT, ipLDDT, ranking score; Metrics where lower is better (right plots): global pAE, and ipAE. **B)** Frequency of each RBD epitope in the PDB, stratified by structures included in the AF3 training set (cutoff 30 September 2021; green) versus structures deposited after the cutoff (red). A structure was assigned to an epitope if ≥2 of its hotspot residues contacted the antibody interface (±2 residues); overlapping epitope definitions allow one PDB entry to contribute to multiple epitopes. **C)** AF3 interface recovery (% correct interface) as a function of epitope representation in the PDB training set, grouped by frequency <50 or >50 structures. **D)** Representative antibody - D614G RBD complex colored by DiscoTope-3.0 epitope-likeness score. **E)** Agreement between AF3 model selection by the default ranking score and by ipSAE (maximum ipSAE across antibody-antigen chain pairs). **F)** Interface recovery for high-affinity (left, green, neutralization IC50 < 1) versus low-affinity (right, pink, 1 < neutralization IC50 < 10) antibodies, stratified by epitope. **G)** Relationship between experimental mutation escape score (single RBD point mutants) and AF3 in silico score delta ipTM (mutated - wild-type, top model over 50 seeds) for a subset of 10 antibodies. The red line shows an ordinary least-squares linear regression with its 95% confidence interval (seaborn regplot); r is the Pearson correlation. A dashed line at ΔipTM = 0 marks no predicted change relative to wild type.

**Figure S6.**
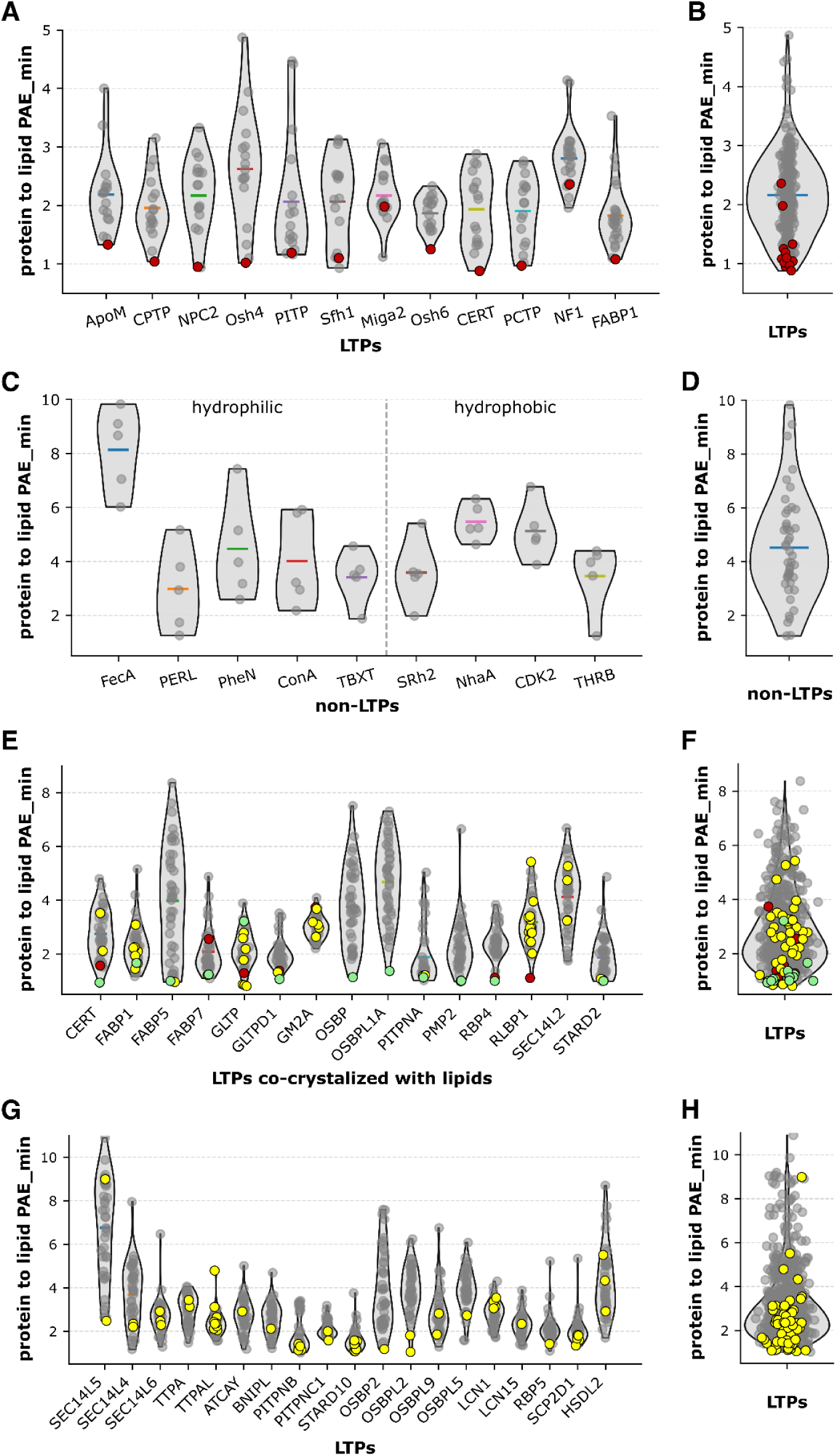
Distribution of protein to lipid minimum interface predicted aligned error (min-iPAE) values obtained from AF3 protein-lipid co-folding predictions across multiple benchmarking datasets. Panels **A)** and **B)** correspond to DB1, **C)** and **D)** to DB2, **E)** and **F)** to DB4, and **G)** and **H)** to DB5. For each dataset, per-protein min-iPAE distributions are shown in the left panels **(A, C, E, and G)**, while aggregated distributions across all proteins are shown in the right panels **(B, D, F, and H)**. Across all panels, violin plots represent the distribution of min-iPAE scores for all predicted protein-lipid pairs. Individual data points corresponding to experimentally validated interactions are overlaid as colored markers: red dots indicate co-crystallized protein-lipid complexes (true positives derived from x-ray structures), yellow dots indicate interactions supported by mass spectrometry or high-performance thin-layer chromatography, and green dots denote interactions validated by both structural and biochemical evidence. All remaining predictions correspond to protein-lipid pairs without experimental support (putative negatives or uncharacterized interactions). The relative positioning of experimentally validated interactions within the min-iPAE distributions illustrates the discriminative performance of AF3 across datasets.

**Figure S7.**
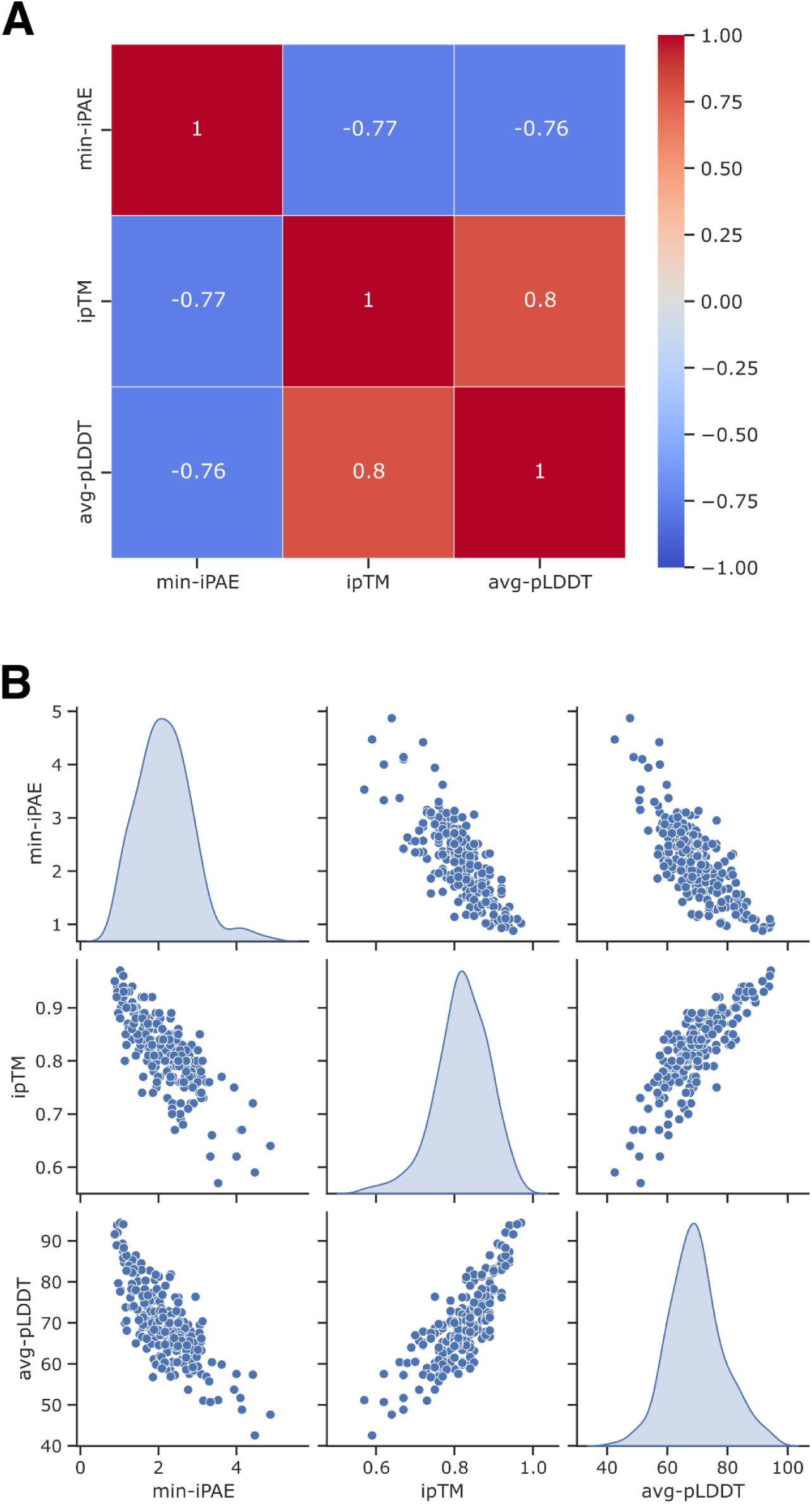
Correlation between AF3 protein-lipid co-folding metrics. **A)** Pearson correlation heatmap and **B)** corresponding scatter plot matrix for the three AF3-derived metrics employed to assess the confidence of the protein-lipid co-folding predictions.

**Figure S8.**
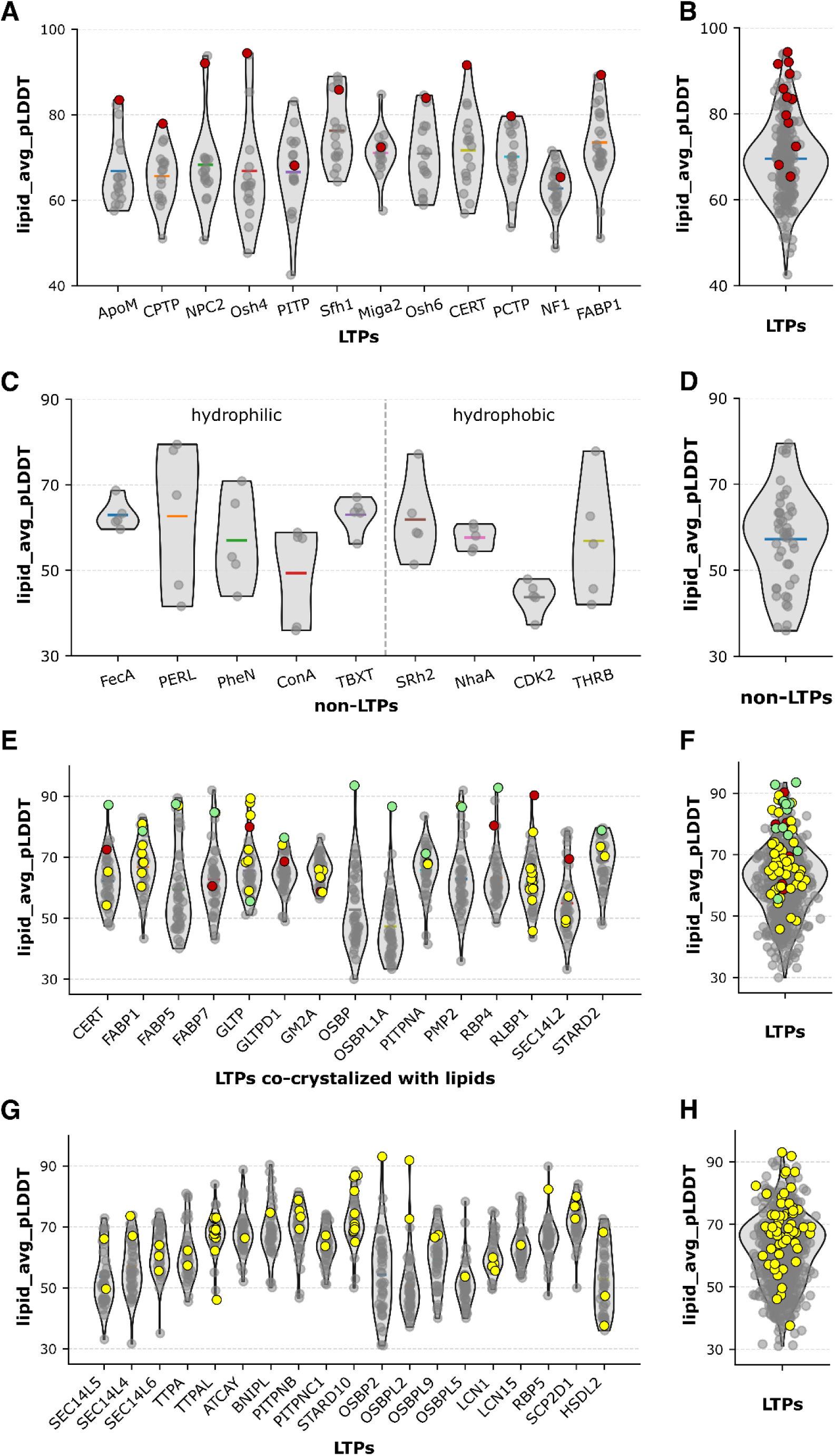
Distribution of average lipid predicted local distance difference test (avg-pLDDT) values obtained from AF3 protein-lipid co-folding predictions across multiple benchmarking datasets. Panels **A)** and **B)** correspond to DB1, **C)** and **D)** to DB2, **E)** and **F)** to DB4, and **G)** and **H)** to DB5. For each dataset, per-protein avg-pLDDT distributions are shown in the left panels **(A, C, E, and G)**, while aggregated distributions across all proteins are shown in the right panels **(B, D, F, and H)**. Across all panels, violin plots represent the distribution of avg-pLDDT scores for all predicted protein-lipid pairs. Individual data points corresponding to experimentally validated interactions are overlaid as colored markers: red dots indicate co-crystallized protein-lipid complexes (true positives derived from x-ray structures), yellow dots indicate interactions supported by mass spectrometry or high-performance thin-layer chromatography, and green dots denote interactions validated by both structural and biochemical evidence. All remaining predictions correspond to protein-lipid pairs without experimental support (putative negatives or uncharacterized interactions). The relative positioning of experimentally validated interactions within the avg-pLDDT distributions illustrates the discriminative performance of AF3 across datasets.

**Figure S9.**
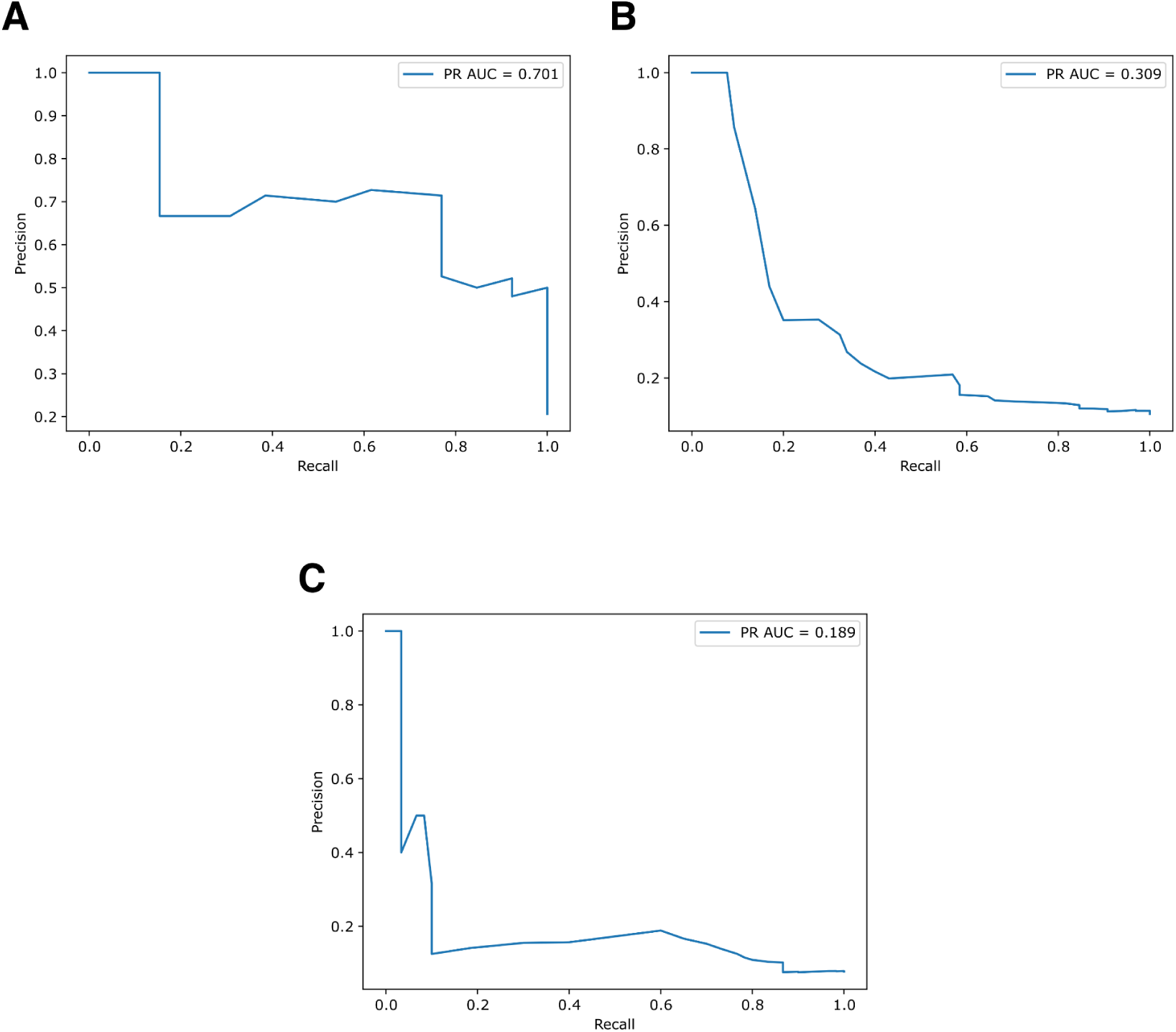
Precision-Recall curves. Panels **A)**, **B)**, and **C)** display the PR curves and the AUC values corresponding to DB3, DB4, and DB5 datasets, respectively.

## List of Supplementary Tables

**Table S1 - Ubiquinated protein structures in the benchmark dataset.** Summary of structures present in the benchmark of ubiquitinated structures.

**Table S2 - Ubiquitin-protein bonds in the benchmark dataset.** Ubiquitin-protein covalent bonds present in the benchmark dataset. Columns beginning with “pdb” show the bonded atom pairs as they exist in the raw pdb file, while columns beginning with “prediction” show the bonded atom pairs as they exist in the processed and predicted structure.

**Table S3 - Results from comparing experimental and predicted ubiquitinated structures.** For each structure, five AlphaFold models were generated with covalent bonds included and five were generated without covalent bonds. Predicted structures were compared to the experimental structure using DockQ and RMSD. Confidence metrics and other values output by AlphaFold3 are included.

**Table S4 - Benchmark dataset and AF3 confidence scores for TCR predictions**. A: Prediction of paired TCR sequences and A0201_LLWNGPMAV epitope. B: Prediction of TCRs from Gumpert et.al for 3 different epitopes. C: Prediction of peptides from YF vaccine and a known TCR. D: Prediction of TCR variants and epitopes. E: Prediction of 3 known TCRs with epitope variants.

**Table S5 - Antibody-antigen complex sequences in the benchmark dataset.** Summary of antibody and SARS-CoV-2 antigen sequences present in the benchmark dataset

**Table S6 - Antibody-antigen complex mutations in the benchmark dataset.** Summary of antibody and SARS-CoV-2 antigen mutations present in the benchmark dataset

**Table S7 - Benchmark dataset for protein-RNA predictions**. List of proteins and RNA sequences used for modelling for different sets of positive and negative pairs.

**Table S8 - Protein-RNA cross-linking data**. Protein-RNA cross-linking position for proteins with known protein-RNA interactions.

**Table S9 - Protein-lipid benchmark datasets**. Compilation of protein and lipid molecules for the 5 sets (DB1-5) compiled for this analysis.

## Notes

### Competing Interest Statement

The authors have declared no competing interest.

